# Changing and stable chromatin accessibility supports transcriptional overhaul during neural stem cell activation

**DOI:** 10.1101/2020.01.24.918664

**Authors:** Sun Y. Maybury-Lewis, Abigail K. Brown, Mitchell Yeary, Anna Sloutskin, Shleshma Dhakal, Brendan McCarthy-Sinclair, Tamar Juven-Gershon, Ashley E. Webb

## Abstract

Adult neural stem cells are largely quiescent, and require transcriptional reprogramming to reenter the cell cycle and undergo neurogenesis. However, the precise mechanisms that underlie the rapid transcriptional overhaul during NSC activation remain undefined. Here, we identify the genome-wide chromatin accessibility differences between primary neural stem and progenitor cells in quiescent and activated states. We show that these distinct cellular states exhibit both shared and unique chromatin profiles, which are both associated with gene regulation. Interestingly, we find that accessible chromatin states specific to quiescent or activated cells are active enhancers bound by pro-neurogenic and quiescence factors, ASCL1 and NFI. In contrast, shared sites are gene promoters harboring constitutively accessible chromatin enriched for particular core promoter elements that are functionally associated with translation and metabolic functions. Together, our findings reveal how accessible chromatin states regulate a transcriptional overhaul and drive the switch between quiescence and proliferation in NSC activation.

## Introduction

Neural stem cells (NSCs) are the source of new neurons, astrocytes, and oligodendrocytes in the adult mammalian brain. Studies in rodents show that adult-born neurons contribute to learning and memory, sensory functions, and mood regulation^1–5^. NSCs are located in distinct niche environments in the brain that provide molecular cues that promote or suppress their growth, proliferation, and differentiation into neurons or glia. There are two well-defined neurogenic niches in the postnatal and adult brain: the subventricular zone (SVZ) lining the lateral ventricles, and the dentate gyrus (DG) of the hippocampus. Similar to rodents and non-human primates, NSCs undergo postnatal neurogenesis in humans, but the exact age at which neurogenesis declines remains controversial^6–9^.

In vivo, the majority of NSCs reside in a state of quiescence^10^. Quiescent NSCs (qNSCs) have exited the cell cycle but can be prompted by intrinsic or extrinsic cues to “activate” and re-enter the cell cycle (we refer to these cells as activated NSCs, or aNSCs). Once activated, aNSCs proliferate and may return to quiescence and self-renew, or differentiate into neurons or glia. Activation of qNSCs is the first critical step in neurogenesis in the adult brain, and is enhanced in response to damage (e.g. stroke) or environmental stimuli such as parabiosis^11–13^. Evidence shows that decreased neurogenesis with age occurs due to reduced activation of qNSCs, senescence of the NSC niche, and exhaustion of the qNSC pool^14–17^. Accumulation of qNSCs has also been observed in a rodent model for neurodevelopmental disorders, suggesting that a careful balance of NSC quiescence and activation is necessary for healthy cognitive function^18^. However, the precise mechanisms that regulate this balance and prompt qNSCs to re-enter the cell cycle in the healthy mammalian brain are mostly unknown.

Recent studies reported that quiescent and activated NSCs employ cell type-specific mechanisms to support their functionality, including distinct metabolic states and differences in proteostasis^19–21^. Transcriptional profiling of qNSCs and aNSCs revealed both shared and distinct transcriptional signatures in the two cell types, indicating that a transcriptional overhaul occurs at a subset of genes during the process of NSC activation^10, 19, 22, 23^. For example, genes involved in cell proliferation, lipid metabolism, and protein homeostasis were differentially expressed in quiescent and activated NSCs.

Similar changes have also been observed using in vitro models of NSC quiescence and activation^24^, which provide an opportunity for more in-depth mechanistic studies. These findings raise the question of how discrete transcriptional states are established across the NSC lineage, and how specific transcriptional changes are regulated at the chromatin level to drive neurogenesis. Understanding these mechanisms will provide important insight into the mechanisms by which intrinsic and extrinsic cues support stem cell functionality in the postnatal and adult brain.

Cell type-specific gene expression is achieved through the use of gene-specific cis regulatory DNA enhancer elements. Enhancers have been extensively studied in the context of development where they are critical for establishing tissue-specific gene expression patterns^25^. In neurodevelopment, for example, gene-specific enhancers that drive expression of key neurogenic regulators such as TLX and DLL1 have been identified^26, 27^. Enhancers typically contain clusters of DNA motifs that function as recognition sequences for specific transcription factors. Through a chromatin looping mechanism, the enhancer regions bound by specific transcriptional regulators physically contact the basal transcriptional machinery at promoter regions to activate or repress transcription^28, 29^. In addition to transcription factor binding, enhancers are characterized by an open chromatin conformation, low nucleosome occupancy, the presence of histone 3 lysine 4 monomethylation (H3K4me1) at flanking nucleosomes, and hypersensitivity to DNA nucleases^30^. Enhancers can be further subdivided into active enhancers marked by acetylated histone H3K27 (H3K27ac) and p300 acetyltransferase binding, and poised enhancers that lack these features^31–33^. Recent studies using mouse neural progenitor cells have used covalent histone marks to define enhancer states in both activated and quiescent cells in vitro, indicating that these elements play a key role in the neurogenic lineage^24, 34^. Distal regulatory enhancers are also critical in human cortical neurogenesis, and have been linked to neuropsychiatric diseases^35–37^.

Here, we used a recently developed assay for chromatin accessibility (ATAC-seq) to map the genomic chromatin states in quiescent and activated NSCs in vitro. We detected regions of chromatin that have distinct accessibility states in quiescent and activated cells, are enriched with marks of active enhancers, and are associated with differential gene expression in vivo in the adult brain. Interestingly, we also observed a number of genomic regions constitutively accessible in both quiescence and activation, yet associated with differential gene expression in vivo. Moreover, genes regulated through active enhancers are critical for supporting the neurogenic lineage, whereas those exclusively associated with constitutively accessible promoters are enriched for metabolic functions and proteostasis. Distal active enhancers were highly enriched for binding of transcriptional regulators of quiescence and activation, NFIX and ASCL1, whereas stable chromatin regions were enriched for CTCF binding, a known insulator. Thus, at least two distinct mechanisms revealed by chromatin accessibility profiling are associated with a reprogramming of the transcriptional circuitry during early neurogenesis. Together, these findings represent the first genomic map linking chromatin accessibility to neurogenic gene regulation in vivo, reveal chromatin states that define the neurogenic lineage, and uncover the key regulatory regions that support activation of quiescent stem cells in the brain.

## Results

### Quiescent and activated NSPCs share a lineage-specific chromatin profile

To investigate how transcriptional changes underlying NSC activation are driven by changes in accessibility of chromatin, we performed ATAC-seq (Assay for Transposase Accessible Chromatin)^38^ on quiescent and activated primary mouse neural stem and progenitor cells (NSPCs; note that we refer to primary NSCs in culture as NSPCs since cultures contain a mixture of stem and progenitor cells^39^). We isolated NSPCs from postnatal mouse brains and collected nuclei from early passage cells cultured in activated/growth conditions or quiescent conditions, as previously established^19, 24, 40^.

Activated NSPCs were maintained in proliferation media containing EGF and FGF2, whereas quiescent NSPCs were cultured in BMP4 (Bone Morphogenetic Protein 4) and FGF2, which induces a rapid, reversible quiescent state in NSPCs^24, 40^. Cell cycle status under the two conditions was confirmed using an EdU (5-ethynyl-2’-deoxyuridine) incorporation assay, and proliferation was significantly reduced in the quiescent cells (*P* < 0.001, Student’s t-test) (Fig. 1a, Supplementary Fig. 1a). To confirm that induced-quiescent NSPCs retain the potential to reenter the cell cycle, we reactivated the BMP4-treated cells by removing BMP4 and adding back the proliferation media containing EGF and FGF2 (Supplementary Fig. 1a-b). We confirmed that activated, quiescent, and reactivated NSPCs expressed the stem cell marker SOX2 (Supplementary Fig. 1c). The activated and reactivated NSPCs expressed higher levels of NESTIN compared to quiescent, consistent with previous findings in the SVZ in vivo^10^. Using this quiescence model system, we generated ATAC-seq libraries from quiescent and activated NSPCs to map the genomic changes in chromatin accessibility in the two cellular states (Fig. 1b). Initial analysis of our ATAC-seq data revealed that quiescent and activated NSPCs have on average over 43,000 accessible chromatin sites, defined by “peaks” in ATAC-seq signals (Supplementary Table 1). We found that ATAC-seq signals between two biological replicates in quiescent and activated conditions were highly correlated (r = 0.94 in quiescent and 0.95 in activated, Pearson correlation coefficient) (Fig. 1c). In contrast, ATAC-seq signals across quiescent and activated replicates were less correlated (r = 0.74 and 0.77 in quiescent versus activated replicates, Pearson correlation coefficient). Next, we used DiffBind to identify the shared and differential ATAC-seq signals between quiescent and activated conditions using biological replicates^41^. Interestingly, comparison of accessible chromatin between quiescent and activated NSPCs revealed that approximately 67% of the accessible sites (19,976 sites) were shared between quiescent and activated conditions. These sites represent stably accessible chromatin regions that do not change significantly between the two cellular states, defining an accessibility signature of the neurogenic stem and progenitors (Fig. 1d). Furthermore, NSPCs exhibit quiescence- and activation-specific open chromatin sites (3,152 and 6,777 sites, respectively) (Fig. 1d). Accessible sites in quiescent and activated NSPCs were similarly distributed genomically, with 42-46% within promoters (-1 kb/+1 kb window around transcription start sites or TSSs), 22-23% in distal intergenic regions, and 25-27% in introns (Fig. 1e). Comparison of quiescent and activated NSPC ATAC-seq signals to those from postnatal mouse lung, liver, intestine, kidney, and stomach tissues (ENCODE)^42^ revealed that NSPCs exhibit a distinct chromatin profile from other cell lineages (Supplementary Fig. 1d). Together, these analyses reveal that quiescent and activated NSPCs exhibit similarities in global chromatin profiles, with quiescence and activation-specific differences in approximately 33% of the accessible chromatin.

**Fig. 1.**
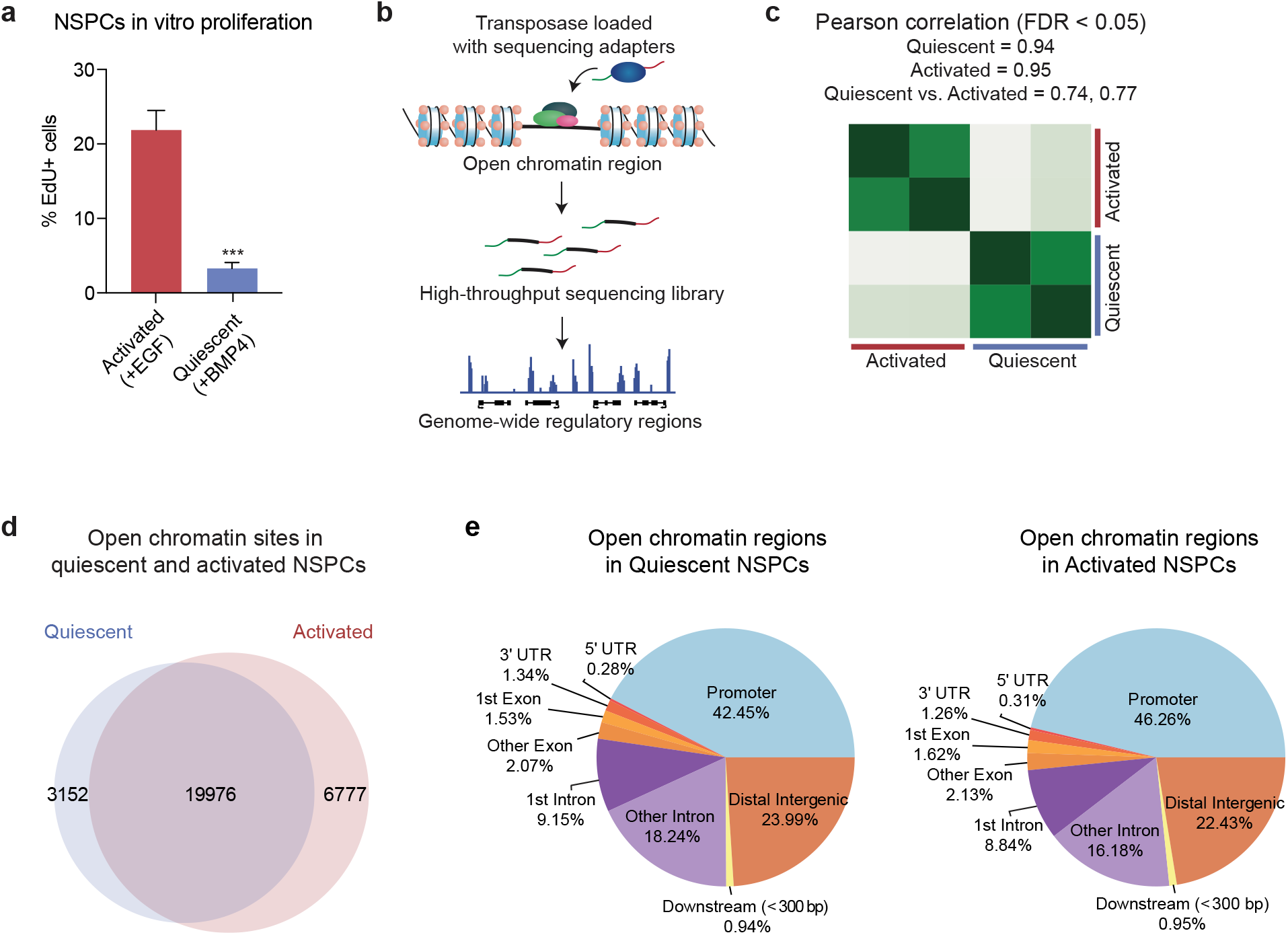
Quiescent and activated NSPCs have shared and unique accessible chromatin regions. (a) EdU incorporation assay to determine cell cycle status of primary activated and quiescent NSPCs used in this study. (n = 3, \*\*\**P* < 0.001, Student’s t-test). (b) Schematic representation of the ATAC-seq assay to detect chromatin accessibility in quiescent and activated NSPCs. (c) Heatmap depicting Pearson correlation coefficient r (0.94 in quiescent replicates, 0.95 in activated replicates) between ATAC-seq biological replicates in quiescent and activated conditions. (d) Venn diagram depicting overlapping and unique accessible chromatin sites in quiescent and activated NSPCs. 19,976 peaks are common across quiescent and activated conditions, 3,152 peaks are quiescence-specific, and 6,777 peaks are activation-specific (FDR < 0.05). (e) Pie charts representing the genomic distribution of all open chromatin regions in quiescent (left) and activated (right) NSPCs. Promoter regions were defined as −1/+1kb around TSSs, and ATAC-seq peaks were annotated by their distance from gene promoters.

### Quiescent and activated NSPCs exhibit distinct states of chromatin accessibility genome-wide

We next examined how chromatin accessibility is altered in specific genomic regions between quiescent and activated NSPCs. As an initial assessment, we investigated two well-known regulators of NSC self-renewal and proliferation: *Olig2* and *Fgf1*^43, 44^. Our ATAC-seq data indicated that in one upstream region associated with *Olig2*, chromatin accessibility was increased in activated NSPCs relative to quiescent (Fig. 2a, top)^42, 45^. We designated such dynamic regions that are significantly more accessible in activated NSPCs compared to quiescent NSPCs as “Accessible in Activated”, or “AA”. In contrast, we observed chromatin accessibility changes in two distinct regions in the first intron of *Fgf1.* Interestingly, in this case, chromatin accessibility was higher in activated NSPCs in one region (Peak 1), and in quiescent NSPCs at a second site (Peak 2) relative to the other cell type (Fig. 2a, bottom). We refer to regions that are significantly more accessible in quiescent NSPCs, such as Peak 2, as “Accessible in Quiescent”, or “AQ”. To examine whether the changes in accessibility correlated with differences in gene expression, we interrogated two independent RNA-seq datasets: 1) freshly isolated quiescent and activated NSCs purified by fluorescence-activated cell sorting (FACS)^19^ and 2) cultured NS5 cells (an embryonic stem cell derived neural progenitor line)^24^. In both datasets, quiescent and activated populations exhibit significant differential expression of *Olig2* and *Fgf1* (Fig. 2b). Furthermore, ChIP-seq datasets for enhancer marks suggest that these dynamic peaks may have functional enhancer activity (Supplementary Fig. 1e).

**Fig. 2.**
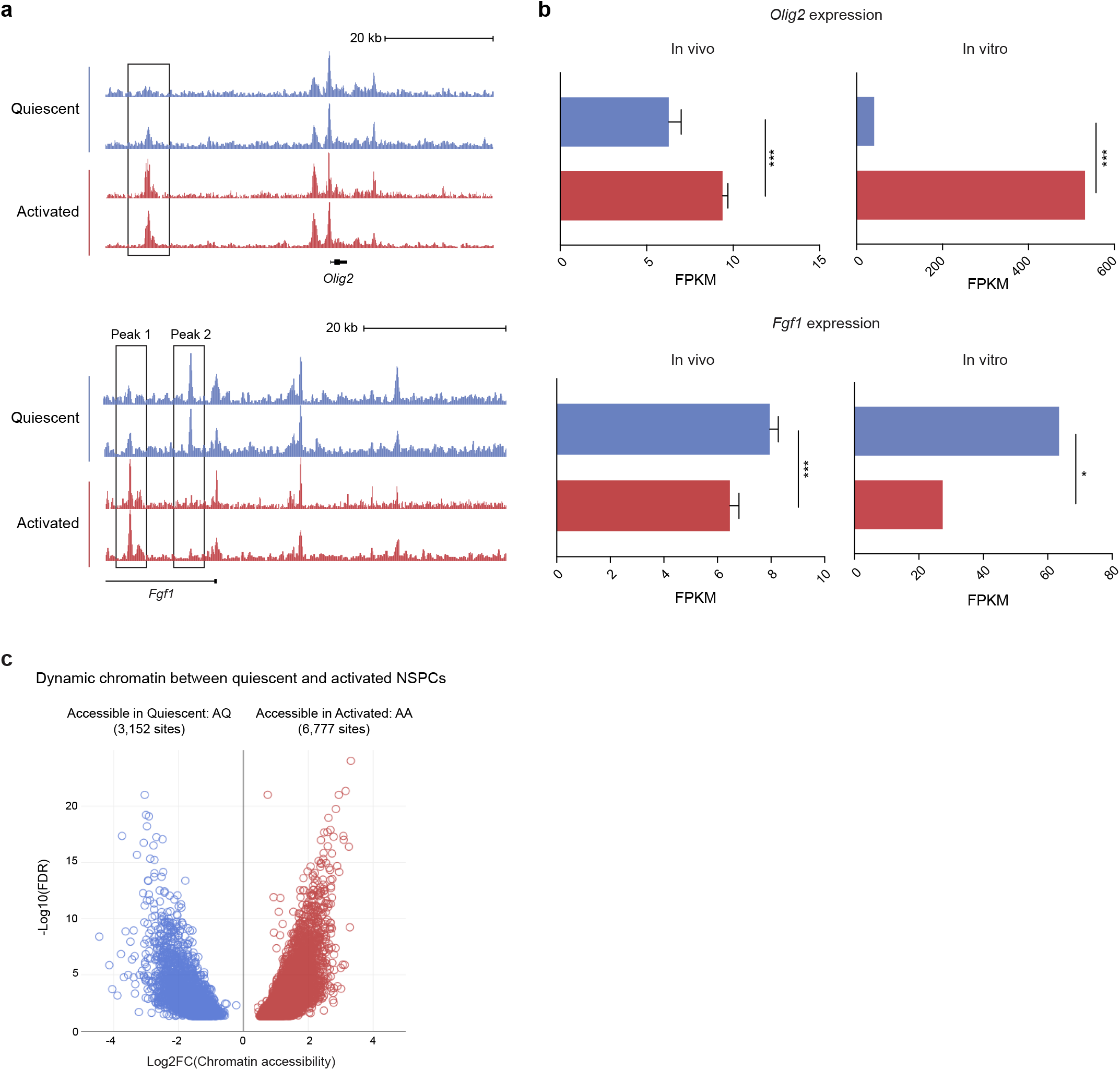
Quiescent and activated NSPCs exhibit differential accessibility in genomic regions. (a) UCSC genome browser shot of representative ATAC-seq tracks at the *Olig2* (top) and *Fgf1* loci (bottom). A region upstream of the *Olig2* TSS (boxed) shows increased accessibility in activated NSPCs compared to quiescent cells. Two regions in the first intron of *Fgf1*, labeled Peak 1 and 2, show increased and decreased accessibility in activated NSPCs compared to quiescent cells, respectively. We refer to peaks such as Peak 1 as “Accessible in Activated”, or “AA”, and peaks such as Peak 2 as “Accessible in Quiescent”, or “AQ”. (b) Bar graphs depicting differential expression of *Olig2* and *Fgf1* in freshly isolated quiescent and activated NSCs in vivo (left), and in NS5 cells in vitro (right). In both datasets, differential expression of *Olig2* and *Fgf1* was statistically significant (FDR-corrected \*\*\**P* < 0.001, \**P* < 0.05). (c) Volcano plot showing fold change in accessibility at all dynamic chromatin sites (AQ + AA) between quiescent and activated NSPCs (FDR < 0.05).

Globally, we examined the genome-wide differences in chromatin profiles of quiescent and activated NSPCs. In total, 9,929 sites had altered accessibility between quiescent and activated NSPCs (FDR < 0.05) (Supplementary Table 2). These sites of dynamic accessibility consisted of 3,152 AQ and 6,777 AA sites (31.8% and 68.2% of all dynamic chromatin, respectively) (Fig. 1d and 2c).

### Chromatin accessibility in reactivated NSPCs largely resembles the activated state

We next compared the global chromatin profiles of reactivated NSPCs to activated and quiescent cells. Comparison of biological replicates in activated, quiescent, and reactivated ATAC-seq revealed that reactivated NSPCs resemble activated cells and are distinct from the quiescent cells (Supplementary Fig. 2a-b). Interestingly, open chromatin regions in the reactivated NSPCs were located less frequently in the promoters (37.14% compared to 42-46% in quiescent and activated), and more often in the distal regions (25.22% in intergenic, 30.73% in intronic regions) (Supplementary Fig. 2c). Stable chromatin regions between quiescent and activated NSPCs were nearly all open in the reactivated (19,766 of 19, 976 sites) (Supplementary Fig. 2d), suggesting that accessibility of these sites is established in quiescence and maintained as open throughout activation. Interestingly, the reactivated cells shared sites specific to both the activated and quiescent cells. But importantly, AA dynamic chromatin regions (6,466 of 6,777 sites) were almost all open in the reactivated cells. In contrast, only 44% of the AQ dynamic chromatin regions (1,393 of 3,152) remained open in the reactivated NSPCs (Supplementary Fig. 2d). Together, these data indicate that the quiescent cells have the ability to revert their chromatin to an activated accessibility state and that the reestablishment of this state is sufficient for complete reentry into the cell cycle.

### Chromatin regions with dynamic accessibility are associated with gene regulation during NSPC activation

To compare gene expression changes between quiescent and activated NSPCs and the chromatin-level changes we observed, we utilized the two independent RNA-seq datasets profiling quiescent and activated NSCs in vivo and cultured NS5 cells in addition to our ATAC-seq data (Fig. 3a). We first asked whether genes that are differentially expressed in quiescence versus activation exhibit altered chromatin accessibility in the two cellular states. We refer to sites with differential accessibility in the two cellular states as AQ/AA, or collectively dynamic sites. In freshly isolated NSCs, 2,088 and 2,255 genes were upregulated and downregulated, respectively, in aNSCs compared to qNSCs (Fig. 3a). We first examined the in vivo gene expression changes with NSC activation with open chromatin, and found that genes with stable or dynamic open chromatin were significantly more upregulated (Fig. 3b, Supplementary Fig. 3a for in vitro expression). We next computed the fractions of upregulated and downregulated genes with associated AA and AQ sites (within the −25 kb/+10 kb window from the TSS) (see Supplementary Table 3 for gene assignments to dynamic chromatin sites). We found that approximately 20% of differentially expressed genes (419 upregulated and 471 downregulated) were associated with dynamic chromatin accessibility. Furthermore, we observed a similar association between differentially expressed genes and chromatin accessibility in cultured NS5 cells (in vivo 20.5% and in vitro 20.0%) (Fig. 3a). Among the genes that are upregulated during NSC activation and contain dynamic sites, 85.0% were associated with chromatin opening (i.e. have at least one accessible AA site) (Fig. 3c, Supplemental Fig. 3b). Thus, a gain in chromatin accessibility is associated with gene activation. Consistent with this notion, 24.4% of genes that are downregulated in activated NSCs relative to quiescent were associated with chromatin closing in the activated state (AQ sites) (Fig. 3c, Supplementary Fig. 3b). These findings are consistent with a model in which increased chromatin accessibility at specific sites allows for recruitment of gene-specific transcription factors or general transcriptional machinery to drive gene expression. Notably, we also observed 69.6% of genes that are downregulated in activated cells compared to quiescent yet contain accessible sites only in the activated state (AA sites). This finding suggests that transcriptional repression is also associated with chromatin remodeling at these target genes.

**Fig. 3.**
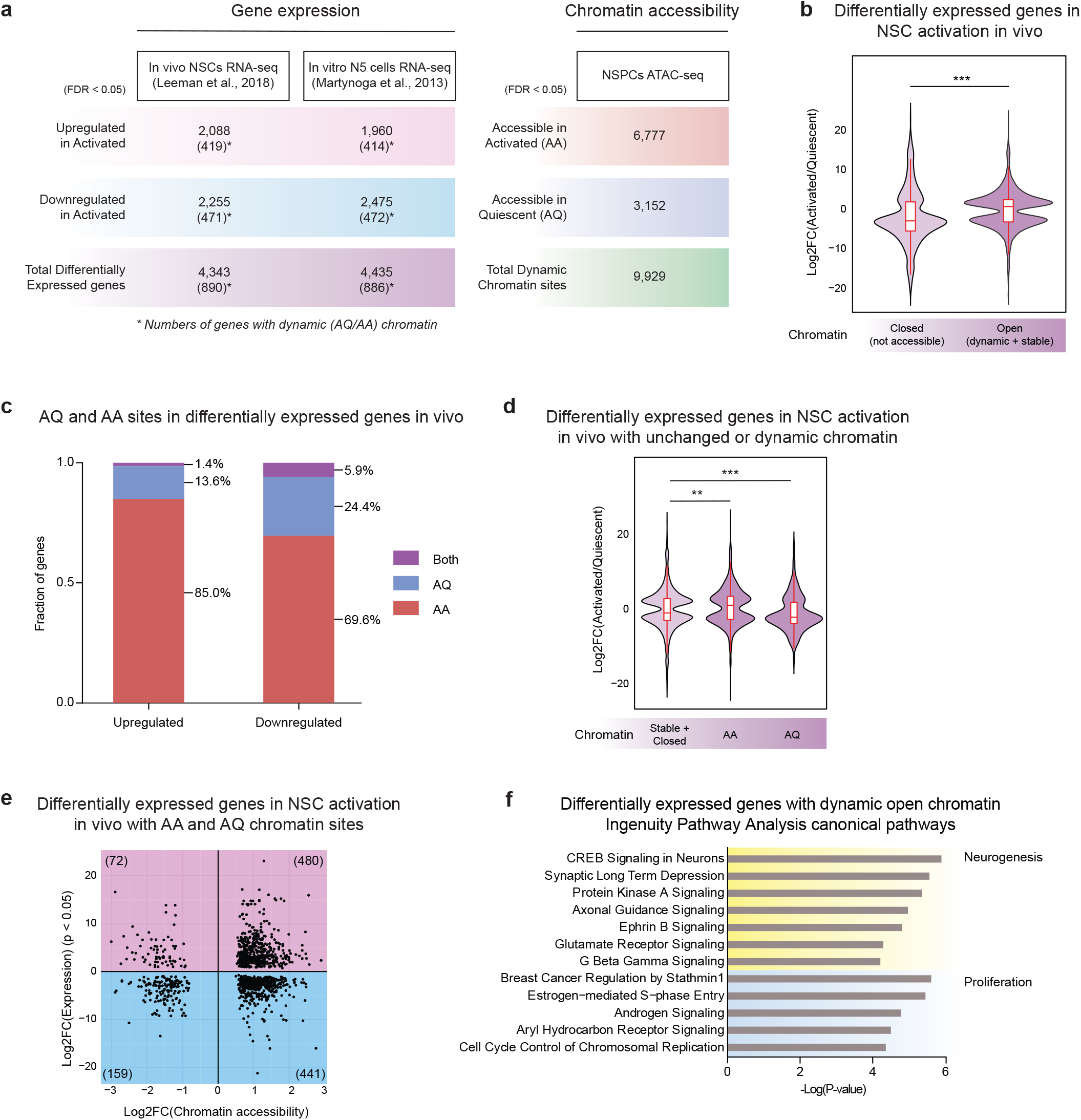
Dynamic chromatin regions are associated with differential expression of genes in quiescent versus activated NSPCs. (a) Summary of the differentially expressed genes in two independently generated RNA-seq datasets and differentially accessible chromatin sites in ATAC-seq (FDR < 0.05). Numbers in parentheses indicate differentially expressed genes with dynamic (AQ or AA) chromatin sites. (b) Comparison of differential expression in vivo (fold change) between genes with closed chromatin (no ATAC-seq signal) and open chromatin (Wilcoxon signed rank test, \*\*\**P* < 0.001). (c) Bar graph depicting percentages of differentially expressed genes in vivo that contain dynamic (AQ or AA) chromatin sites, or both. (d) Comparison of differential expression in vivo (fold change) between genes with chromatin that does not change, AA chromatin, and AQ chromatin (Wilcoxon signed rank test, \*\*\**P* < 0.001, \*\**P* < 0.01). (e) Scatterplot showing fold change in chromatin accessibility versus fold change in associated gene expression (FDR < 0.05). Each dot represents a dynamic chromatin site that is associated with a differentially expressed gene, and the total number of sites in each quadrant is shown in parentheses. Downregulated genes are enriched with AQ sites (Fisher’s exact test, *P* = 2.6 × 10^−7^). (f) Ingenuity Pathway Analysis (IPA) Canonical Pathways of genes differentially expressed in vivo and associated with dynamic chromatin sites (AQ or AA). Top 12 pathways are shown. Complete table of results is shown in Supplementary Table 5.

We next asked whether genes with accessible chromatin states specifically in quiescent or activated cells (AQ or AA sites) have greater gene expression changes compared to those without dynamic chromatin sites. Our analysis revealed that genes with AQ or AA sites were more differentially expressed between quiescent and activated cells compared to genes with no changes in accessibility (Fig. 3d). For example, genes with AA sites exhibited increased expression in activated cells compared to genes without chromatin changes. In contrast, genes with AQ sites displayed increased expression in quiescent cells compared to genes with stable chromatin. Interestingly, genes with AA sites were similarly distributed in up- and downregulated expression in activated NSPCs, while genes with AQ sites were largely downregulated in the activated cells compared to quiescent cells (compare distributions in AA and AQ in Fig. 3d and Supplementary Fig. 3c). Specifically, 480 upregulated and 441 downregulated genes with NSC activation in vivo harbored AA sites; in contrast, 72 upregulated and 159 downregulated genes had AQ sites (Fig. 3e, Supplementary Fig. 3d for in vitro expression). In other words, we found that increased chromatin accessibility is associated with both transcriptional upregulation and repression, whereas chromatin closing is predominantly associated with downregulation of gene expression. Notably, nearly all of the AA chromatin regions associated with up- and downregulation of gene expression became accessible again in the reactivated NSPCs, further supporting our findings (Supplementary Fig. 2d-e). To investigate the functional relevance of changes in chromatin accessibility, we tested whether the genes associated with dynamic chromatin (AQ and AA) sites were enriched for particular signaling pathways using Ingenuity Pathway Analysis (IPA)^46^ (Fig. 3f).

Interestingly, genes with dynamic chromatin were most highly enriched with neural identity, differentiation, and proliferation pathways. Together, our findings suggest that chromatin-level and transcriptomic changes during NSPC activation are linked, and occur at key genes regulating proliferation and differentiation in the neural lineage.

### Constitutively accessible sites are enriched for H3K4me3 in quiescent and activated cells

Our observation that many chromatin regions are accessible in both the quiescent and activated states (stable chromatin sites) raised the question of how these sites function mechanistically to support the NSC lineage. The chromatin regions with stable accessibility were associated with 12,779 genes (Supplementary Table 3), 10,683 of which were exclusively associated with stable chromatin (no dynamic chromatin).

Moreover, genes with only stable open chromatin were associated with upregulation of expression in the activated condition compared to quiescent both in vivo (*P* = 9.78 × 10^−30^, Wilcoxon signed rank test) and in vitro (*P =* 7.63 × 10^−5^, Wilcoxon signed rank test). Our finding that both dynamic (AQ and AA) chromatin and stable chromatin states could be linked to transcriptional activation suggests that differential expression between quiescent and activated NSCs is supported by at least two distinct mechanisms.

Consistent with this possibility, we observed that dynamic and stable chromatin sites had different distributions relative to transcription start sites (Fig. 4a). 57.08% of stable chromatin regions were in promoters (-1 kb/+1 kb from TSSs), and 18.18% were distal intergenic (between genes and outside the promoter window), with another 19.22% in intronic regions. In contrast, chromatin regions with differential accessibility in quiescent and activated NSPCs were more frequent in distal intergenic (33.42%) or intronic (40.45%) regions, and resided less frequently in promoter regions (18.09%). These findings indicate that the major differences in chromatin accessibility in quiescent versus activated NSPCs occur at sites away from promoters, possibly at gene-specific regulatory elements. In addition, our observation that many promoter regions are in a readily accessible state in the quiescent NSPCs, and remain open in activated cells, suggests that accessibility at promoters may be critical to allow rapid toggling between quiescence and activation without a need for chromatin remodeling.

**Fig. 4.**
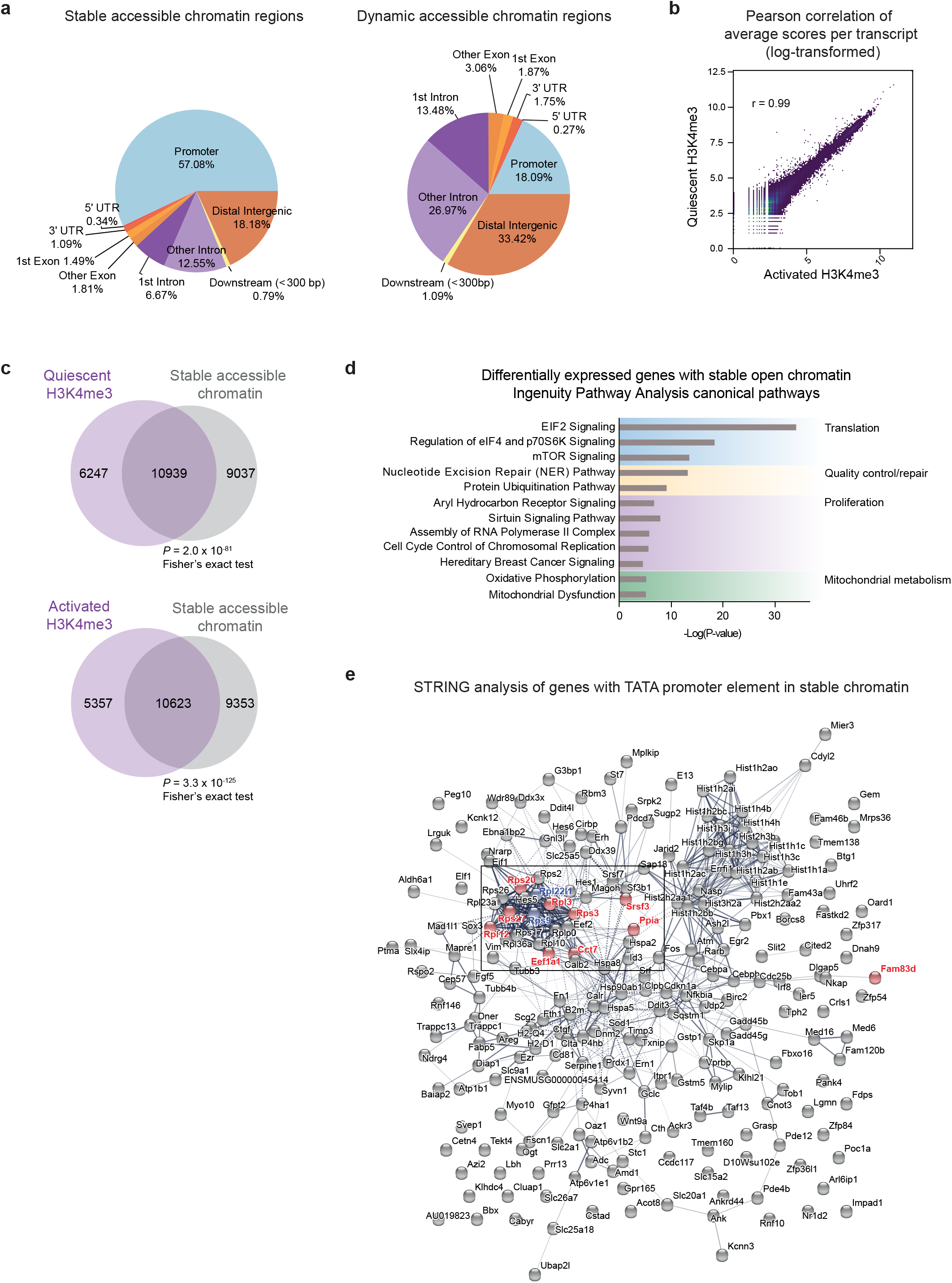
Constitutively accessible chromatin regions in NSPCs are enriched with active promoter marks and are functionally distinct from dynamic chromatin regions. (a) Genomic distribution of chromatin regions that are stably open in quiescent and activated NSPCs (left) and regions that are dynamic (AQ or AA) (right). (b) Scatterplot depicting the Pearson correlation of average H3K4me3 ChIP-seq MACS enrichment scores per gene between quiescent and activated NSPCs (r = 0.99). (c) Overlaps between stably open chromatin sites and H3K4me3-enriched sites in quiescent NSPCs (top) (Fisher’s exact test, *P* = 2.0 × 10^−81^) and activated NSPCs (bottom) (Fisher’s exact test, *P* = 3.3 × 10^−125^). (d) Ingenuity Pathway Analysis (IPA) Canonical Pathways enriched in genesets that are differentially expressed between quiescent and activated NSCs in vivo and harbor only constitutively open chromatin. Top 12 pathways are shown. Complete table of results is shown in Supplementary Table 5. (e) STRING analysis of all genes with the TATA box motif in constitutively accessible promoters. Genes upregulated in activated NSCs in vivo compared to quiescent cells with the TATA box motif within biologically relevant positions (-40/-41 to −13/-14 bp from TSS) are shown in red. Genes containing both the TATA box and TCT motifs (-11 to +16 bp from TSS) in their promoters are shown in blue. The top quartile of upregulated genes are clustered together, consisting of ribosomal and translation-related genes (boxed). Complete lists of the top quartile upregulated genes with the TATA box or TCT motifs are shown in Supplementary Table 6.

Based on our finding that stable chromatin is associated with differential gene expression and found at promoter regions, we examined whether these regions are enriched with H3K4me3, a histone mark for active promoters. We performed H3K4me3 ChIP-seq in quiescent and activated NSPCs, and found that H3K4me3 profiles were highly similar between the two states (r = 0.99, Pearson correlation coefficient) (Fig. 4b). Furthermore, we found that the H3K4me3 signals in NSPCs were distinct from those of various adult and embryonic mouse tissues^47^, defining the neurogenic lineage profile (Supplementary Fig. 4a-b). In total, we identified 17,186 H3K4me3 peaks in quiescent NSPCs, and 15,980 peaks in activated NSPCs (FDR < 0.05) (Supplementary Table 4). Interestingly, 63.65% of quiescent H3K4me3 peaks and 66.48% of activated H3K4me3 peaks were in stable chromatin, indicating that chromatin readily accessible prior to NSC activation is enriched with stable H3K4me3 at the promoters of active genes in the NSC lineage (*P* = 2.0 × 10^−81^ for quiescent and 3.3 × 10^−125^ for activated, Fisher’s exact test) (Fig. 4c). Furthermore, we found that differentially expressed genes are primed with H3K4me3 in their promoters. Out of 4,313 differentially expressed genes between in vivo quiescent and activated NSCs, 73% (3,165 genes) were marked by H3K4me3 in both states. These findings suggest that H3K4me3 is stable in promoters of genes that are differentially expressed with NSC activation, consistent with its enrichment in constitutively accessible chromatin.

Next, we performed enrichment analysis using IPA to interrogate the pathways regulating the genes with stable chromatin. Interestingly, genes differentially expressed with in vivo NSC activation and exclusively harboring stable chromatin (no associated dynamic sites) were enriched for canonical pathways regulating translation, proteostasis, metabolism, RNA polymerase II assembly, and proliferation (Fig. 4d, Supplementary Table 5), in contrast with high enrichment of neural identity, differentiation, and proliferation in dynamic chromatin (Fig. 3f, Supplementary Table 5). In order to better understand how genes lacking dynamic chromatin sites undergo transcriptional regulation, we performed an enrichment analysis of core promoter elements in genes upregulated with NSPC activation using ElemeNT^48^. Analysis of upregulated genes in the top quartile (differential expression FDR < 1.44 × 10^−7^) revealed that they were enriched with the TATA box or human TCT elements (*P =* 1.14 × 10^−5^ and 1.14 × 10^−4^ respectively, hypergeometric analysis) (Supplementary Table 6). Intriguingly, the top quartile of upregulated genes with these promoters are closely associated in a “translation cluster” comprising ribosomal proteins, consistent with the enrichment of translation and proteostasis pathways in the stable open chromatin and with the enrichment of the TCT motif in promoters of ribosomal protein genes and translation initiation and elongation factors (Fig. 4d-e, Supplementary Fig. 4c, Supplementary Table 6). Thus, specific promoter elements at constitutively accessible promoters are likely involved in fine tuning the activity of the transcriptional machinery to properly support changes in basal metabolic states as stem cells activate.

### A subset of distal chromatin regions with changing accessibility during NSPC activation are active enhancers

The majority of the dynamic chromatin sites we identified between quiescent and activated cells (73.87%) reside in distal intergenic and intronic regions (Fig. 4a), suggesting that they may be functional enhancers. We next investigated whether distal AQ/AA sites are enriched for epigenomic marks of functional enhancers. We analyzed markers of active enhancers (H3K27ac and p300 ChIP-seq signal) from a neural progenitor line (NS5 cells) either in actively dividing cells or cells that have been induced to enter quiescence^24^. We compared average enrichment levels for these chromatin features in quiescent cells and activated cells specifically at dynamic chromatin sites. We found a strong enrichment for active enhancer marks from quiescent cells at AQ sites, and from activated cells in AA sites (Fig. 5a). Furthermore, these active enhancer marks were enriched in distal intergenic AQ and AA sites (Fig. 5b), confirming that distal chromatin regions with dynamic accessibility harbor quiescent and activated neural stem lineage-specific active enhancers.

**Fig. 5.**
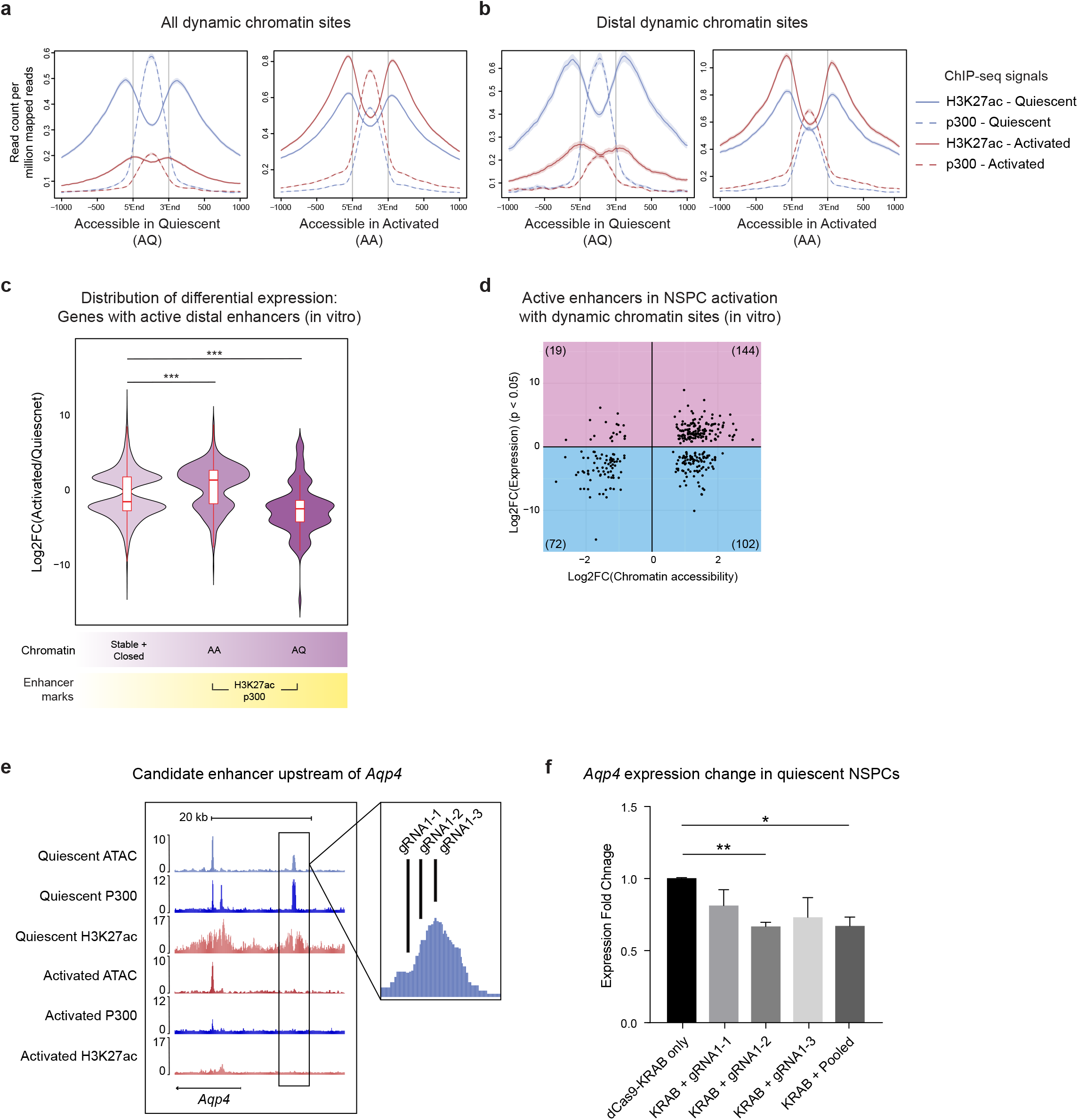
Dynamic chromatin regions are enriched for features of active enhancers and are associated with differential gene expression in NSPC activation. (a) Dynamic chromatin sites (AQ or AA) are enriched for features of active enhancers (H3K27ac and p300). H3K27ac and p300 ChIP-seq signals in quiescent and activated NSPCs in AQ chromatin regions (left) and in AA chromatin regions (right) were plotted in 5’ to 3’ direction. (b) H3K27ac and p300 ChIP-seq signals in quiescent and activated NSPCs, only in distal AQ chromatin (left) and distal AA chromatin (right). (c) Comparison of differential expression of genes with chromatin showing no change (including constitutively open and closed chromatin), and dynamic chromatin (AA or AQ) with active enhancer marks (Wilcoxon signed rank test, \*\*\**P* < 0.001). (d) Scatterplot depicting fold change in chromatin accessibility at active enhancers versus fold change in associated gene expression in vitro (FDR < 0.05). Each dot represents a dynamic site with active enhancer marks that is associated with a differentially expressed gene, and the total number of sites in each quadrant is shown in parentheses. (e) UCSC genome browser snapshot of the putative enhancer region upstream of *Aqp4* locus. gRNAs targeting the enhancer region is shown in a zoomed-in view (right). (f) RT-qPCR analysis showing downregulation of *Aqp4* by dCas9-KRAB with three different gRNA targeting the enhancer region. Fold change for *Aqp4* expression is relative to the dCas9-KRAB only control (n = 3 experiments, Student’s t-test, \*\**P* < 0.01, \**P* < 0.05).

Next, we asked whether changes in chromatin accessibility at these active enhancers are correlated with transcriptional regulation of their associated genes. We found that genes with active enhancers in distal AA sites were more highly expressed on average in activated NSCs compared to quiescent in the RNA-seq dataset from neural progenitor line (NS5 cells) (Fig. 5c). We observed a similar trend between active enhancers in distal AA sites and in vivo gene expression, although it did not reach statistical significance. We suspect this is because chromatin opening at active enhancers is associated with both up and downregulation of gene expression (Fig. 5c-d, Supplementary Fig. 5a-b), consistent with our finding in global chromatin opening (Fig. 3d). Furthermore, active enhancers in distal AQ sites were significantly associated with decreased gene expression in vivo and in vitro (Fig. 5c-d, Supplementary Fig. 5a-b).

We next asked whether targeting enhancers in dynamic chromatin was sufficient to regulate gene expression. We identified an upstream enhancer region (chr18:15421139-15421834) near the *Aqp4* locus. *Aqp4* is expressed in adult NSCs and astrocytes, and has been shown to function in NSC proliferation, survival, migration, and differentiation^49^. Notably, the putative enhancer region we identified was accessible only in quiescent NSPCs and enriched with active enhancer marks from quiescent cells (Fig. 5e). Based on the in vivo and in vitro expression data (Supplementary Fig. 5c), we used the deactivated Cas9 fusion with transcriptional repressor domain KRAB (dCas9-KRAB)^50^ to target the *Aqp4* enhancer in quiescent NSPCs (Supplementary Fig. 5d). We found that one of the three gRNAs targeting this region in dCas9-KRAB-expressing, quiescent NSPCs was sufficient to downregulate *Aqp4* expression (*P* = 0.0024, Student’s t-test) (Fig. 5f). These results demonstrate that enhancers residing in dynamic chromatin function in gene regulation, and targeting these distal elements can modulate gene expression.

Together, our data show that chromatin opening and closing in active enhancers and stable accessibility in promoters are significantly correlated with differential gene expression during NSPC activation. The bimodal distribution of fold change in expression associated with distal AA sites (Fig. 5c-d) indicates that accessible chromatin at active enhancers is associated with both transcriptional activation and repression in the active state relative to quiescence. Supporting this notion, we observed that active enhancers with chromatin opening (distal AA) were associated with upregulation of genes that function in cell cycle and proliferation. In contrast, downregulation was associated with cell signaling pathways such as retinoic acid and Wnt signaling (Supplementary Fig. 5e). Together, these results suggest that active enhancers that become accessible as cells transition from quiescence to activation may be used to both repress and activate gene expression, whereas AQ enhancers primarily function to promote gene expression in the quiescent state.

### Dynamic chromatin regions are bound by major regulators of NSC quiescence and proliferation

Our findings suggest that alterations in chromatin accessibility at specific genomic regions drive transcriptional changes during NSC activation through an enhancer-driven program. We next sought to identify the specific transcription factors that bind accessible chromatin in NSPCs to regulate gene expression. We first performed *in silico* motif analyses of the AQ and AA sites to identify specific transcription factor consensus sequences enriched in these regions. We found that a number of binding motifs were enriched in the dynamic chromatin sites, such as Homeobox, Nuclear Receptor (NR), SMAD, High-mobility group (HMG), and Zinc finger (Zf) motifs (Fig. 6a). Interestingly, AQ and AA sites harbored distinct transcription factor motif sequences. For example, AQ sites were most highly enriched with Nuclear Factor-I (NFI/CTF) motifs (*P* = 1 × 10^−113^), whereas AA sites were most highly enriched for basic helix-loop-helix (bHLH) motifs (*P* = 1 × 10^−-372^). Surprisingly, stably accessible chromatin was most highly enriched for binding motifs of CTCF (*P =* 1 × 10^−1169^), Sp1/5 (*P* = 1 × 10^−316^) and other transcription factors that are known to regulate stem cell proliferation, such as Nuclear Factor Y (NFY) and ETS (*P* = 1 × 10^−315^ and 1 × 10^−248^, respectively)^51–53^. Together, our findings indicate that cell type-specific families of transcriptional regulators are likely responsible for maintaining NSPCs in a quiescent versus activated state through distinct enhancer networks. Moreover, a third discrete set of factors likely bind a transcriptionally competent or stable network of promoter sites (e.g. Sp1) or insulator regions (e.g. CTCF) early on in the neurogenic lineage and also function at these sites as the cells activate.

**Fig. 6.**
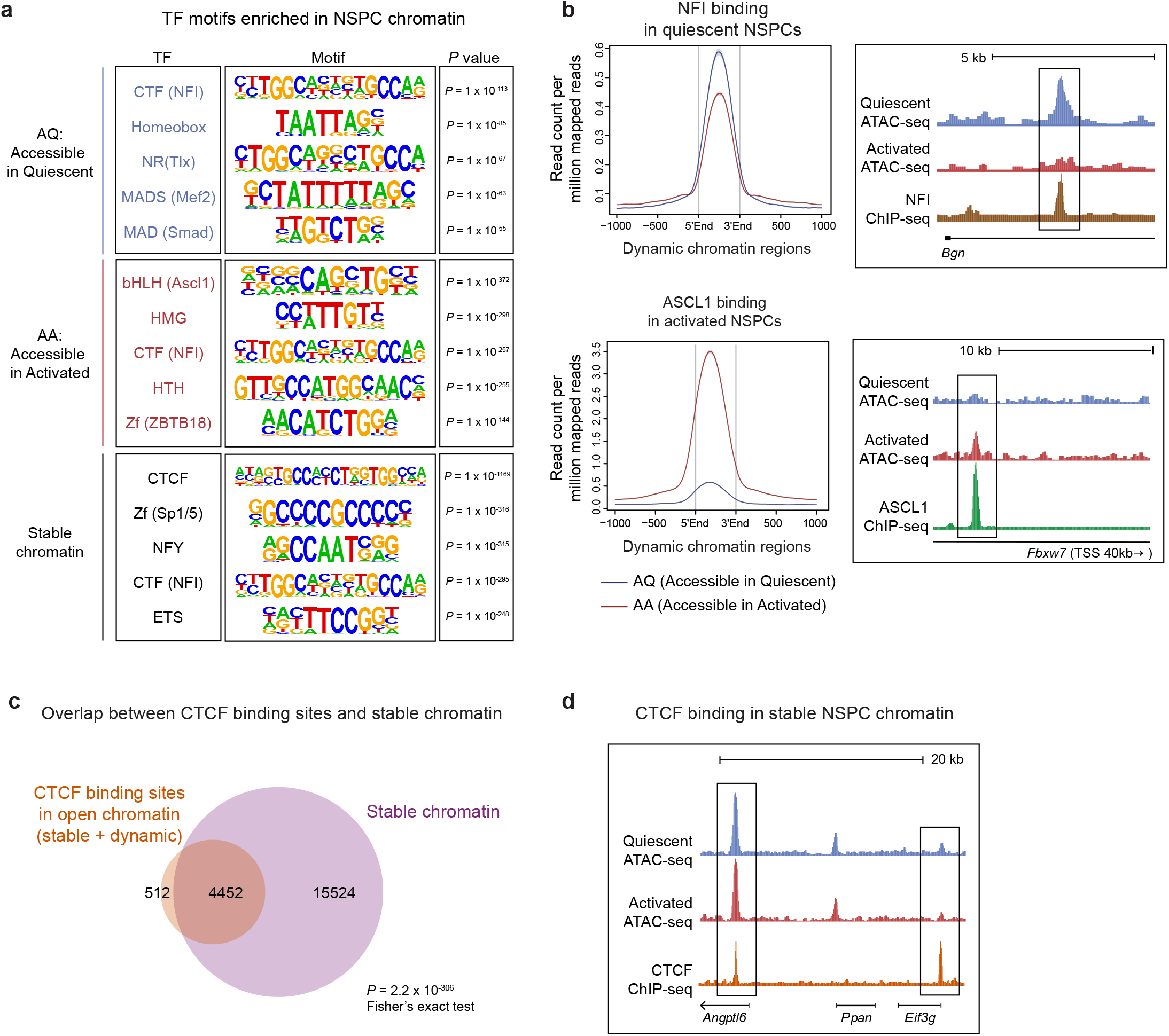
Dynamic chromatin sites are bound by NFI and ASCL1, major regulators of NSPCs, while stable chromatin sites are enriched for CTCF. (a) Summary of Homer motif analysis in AQ, AA, and constitutively accessible chromatin regions. The top 5 most highly enriched motifs are shown for each chromatin state. (b) Pan-NFI (top) and ASCL1 (bottom) ChIP-seq signals from quiescent and activated NSPCs plotted in 5’ to 3’ direction in dynamic chromatin regions (left). UCSC genome browser snapshots of a representative AQ chromatin site with NFI binding (top) and an example of an AA chromatin site with ASCL1 binding (bottom) are shown on the right. (c) Overlaps between constitutively accessible chromatin and CTCF binding open chromatin sites (Fisher’s exact test, *P* = 2.2 × 10^−306^). (d) UCSC genome browser snapshot examples of constitutively accessible chromatin that binds CTCF.

We next investigated the binding distribution of the transcriptional regulators that were the top hits in the motif analysis at AQ and AA site. The most highly enriched motif in the AQ sites was a binding site for the NFI (Nuclear Factor I) transcription factors. NFIX, a member of the NFI transcription factor family, has previously been identified as a major regulator of NSC quiescence in vitro through binding at enhancers^24^. In contrast, AA sites were most highly enriched for binding sites recognized by basic-helix-loop-helix (bHLH) transcription factors. In previous studies, a particular bHLH family member, ASCL1, has been shown to be critical for NSC proliferation and neurogenesis^54–56^. To test whether the AQ and AA sites are primarily bound by these factors in the quiescent and activated states, we overlapped the AA and AQ sites with previously published ASCL1 and pan-NFI ChIP-seq datasets^24, 34^. NFI binding was assayed in quiescent (BMP4-treated) NSPCs, and ASCL1 binding in activated NSPCs, closely recapitulating the conditions under which NSPC chromatin accessibility was examined. Real time quantitative PCR (RT-qPCR) confirmed enrichment for *Nfix* in quiescent NSPCs (Supplementary Fig. 6a-b). *Ascl1* transcript was detectable by RT-qPCR in both quiescent and activated NSPCs (Supplementary Fig. 6a), consistent with previous findings^57^, but enriched in aNSCs RNA-seq in vivo. Globally, we observed that NFI binding was more enriched in AQ sites compared to AA sites (*P* = 2.1 × 10^−29^, Fisher’s exact test), and ASCL1 binding was more enriched in AA sites compared to AQ sites (*P* = 2.2 × 10^−306^, Fisher’s exact test) (Fig. 6b and Supplementary Fig. 6c-d). Specific examples of NFI and ASCL1 binding at AQ and AA sites in the *Bgn* and *Fbwx7* genes are shown in Fig. 6b.

CTCF is a known insulator protein associated with boundaries of topologically associated domains (TADs) of chromatin, functionally preventing inappropriate long-range interactions between promoters and enhancers outside of the TAD^58–60^. Based on our motif analysis, we examined the extent to which CTCF binding is enriched in stable open NSPC chromatin. We analyzed publicly available CTCF ChIP-seq data from activated NSPCs and determined the overlap between these data and our accessible regions^61^. We found that CTCF binding was significantly enriched in the stable chromatin compared to dynamic chromatin (*P* = 2.2 × 10^−306^, Fisher’s exact test) (Fig. 6c-d). Thus, a subset of stable sites are occupied by CTCF, which is likely important for maintaining the higher order chromatin organization during neurogenesis.

Taken together, our findings on enrichment of ASCL1 binding in AA sites, NFI binding in AQ sites, in addition to the motif analyses, strengthen our findings that chromatin sites of dynamic accessibility between quiescent and activated states contain important regulatory elements for NSC activation. Furthermore, enrichment of CTCF motif and binding suggest that TAD boundaries are present in open chromatin that does not change in NSPC activation. These findings support a model in which stable and dynamic regulation of chromatin accessibility is one of the mechanisms by which transcriptional changes are modulated during NSC activation.

## Discussion

We used ATAC-seq to map the chromatin accessibility landscape of primary NSPCs in states of quiescence versus proliferation to uncover the chromatin-level mechanisms supporting the first stage of mammalian neurogenesis. Using RNA-seq expression datasets generated from in vitro and in vivo NSCs, we dissected the relationship between alterations in chromatin accessibility and transcriptional overhaul in NSC activation. The in vitro quiescence system utilized in this study has been shown to be a reliable tool to study the transcriptional profiles of in vivo NSCs^24, 57, 62^. Despite some differences between the datasets, we found strong correlations between chromatin accessibility and gene expression changes in vivo and in vitro. We observed that active enhancers of differentially expressed genes exhibit dynamic accessibility between the quiescent and activated states and are bound by major neural regulators, ASCL1 and NFI. Furthermore, we discovered that approximately 67% of the open chromatin regions are shared between quiescent and activated NSPCs, indicating that at many sites, accessibility is established early in the neurogenic lineage, but can still be associated with gene regulation during stem cell activation. Consistent with this finding, we found that stable chromatin is largely located at gene promoters, enriched for stable H3K4me3, and putative chromatin boundaries bound by CTCF. Furthermore, stable chromatin at core promoters of the top quartile of upregulated genes is enriched with the TATA box and/or TCT motifs and supports coordinated upregulation of ribosomal genes with NSC activation.

In dynamic chromatin, new sites for transcription factor binding that were previously occluded in the quiescent state are exposed with NSC activation, while other sites become less accessible. Our results indicate that these dynamic chromatin regions regulate the balance and transition between NSC quiescence and activation through an enhancer-driven program. Enhancer activity in dynamic chromatin during NSC activation is likely coordinated by alterations in chromatin accessibility, activity of transcriptional co-activators such as p300 to modify histones, and binding of transcription factors for long-range communication with promoters for gene regulation. At active enhancers, both chromatin opening and closing significantly correlate with gene regulation. Unexpectedly, increased chromatin accessibility at enhancers was associated with both up and downregulation of gene expression, while decreased accessibility at enhancers was predominantly associated with downregulation of genes. Moreover, enhancers in chromatin opening with NSC activation are enriched with distinct upregulated and downregulated pathways, suggesting that enhancers are likely bound by both activators and repressors and their function depends on the balance between them. These observations implicate enhancer-driven programs in the transcriptional overhaul required for switching between active and dormant cellular states. While the molecular mechanisms preceding and driving chromatin opening and closing at active enhancers in NSC activation remain unknown, our work has for the first time identified the active enhancers with dynamic chromatin accessibility and their relationship to changes in gene expression. Moreover, we have shown that repressing an active enhancer region is sufficient to induce changes in target gene expression. Currently, identifying functional target genes of enhancers genome-wide remains a challenging task, as enhancers are thought to target multiple promoters for gene regulation^30, 63^. Emerging evidence shows that chromatin accessibility and presence of the H3K27ac histone mark are strong indicators of enhancer activity for cell type-specific genes^64^. While attributing individual enhancer activity to differential expression of their target genes may be difficult, we have found that collectively, dynamic accessibility of the enhancers during NSC activation is significantly associated with gene expression changes nearby. In future work, incorporating information of 3-dimensional chromatin conformation at higher resolution than current limitations will be important to more accurately relate the local chromatin changes revealed in this study to regulation of specific genes.

Using a combination of in silico motif mining and ChIP-seq analysis, we found that the transcriptional regulators NFI and ASCL1 each bind many chromatin regions that are uniquely accessible in quiescent or activated cells, respectively. Previous studies implicated these factors as key regulators of neurogenesis in vivo and in vitro^24, 54–56, 65^. Our observation raises an important question about the relationship between chromatin alterations and transcription factor binding during stem cell activation. Interestingly, ASCL1 was recently identified as a “pioneer factor” that can establish cell type-specific chromatin states in the context of cellular reprogramming. Pioneer factors initiate transcription of silent genes by binding closed chromatin and establishing accessibility for other transcription factors during development and cell reprogramming^66, 67^. In the case of ASCL1, its binding is enriched in chromatin regions that gain accessibility and nucleosome phasing as fibroblasts are directly reprogrammed to induced neurons (iNs)^68^. Therefore, our findings that ASCL1 binds chromatin regions that become more accessible in the activated cells (AA sites) raise the exciting possibility that ASCL1 functions as a pioneer factor during NSC activation as well. Similarly, our findings suggest that NFI factors may also function as pioneer factors to reset chromatin as cells reenter quiescence. Moreover, while NFI motif was most highly enriched in AQ chromatin, its enrichment in AA and stable chromatin suggests that NFI factors are multifunctional in the neurogenic lineage. Supporting this finding, NFIX has been shown to function in neuroblast differentiation in the mouse hippocampus^65^. In future work, it will be important to identify the precise sequence of transcription factor binding, chromatin remodeling events, and transcriptional alterations that occur as quiescent NSCs enter the cell cycle and return to quiescence in order to fully understand how transcriptional programs are regulated to maintain the balance between NSC quiescence and activation. A number of other transcriptional regulators have been shown to regulate the neurogenic lineage, including FOXOs, TLX, DLX, and SOX2, and it will be useful to incorporate data on these transcriptional networks as it becomes available to gain an integrated understanding of how neurogenic transcriptional programs are established and maintained^26, 34, 69–73^. Moreover, it will be critical to determine the mechanism by which ASCL1 functions as a pioneer factor during neurogenesis, either through acting alone or through the recruitment of ATP-dependent chromatin remodeling factors.

Apart from the dynamic chromatin regions, we observed that approximately 67% of the open chromatin regions in NSPCs are accessible in both quiescent and activated states, which we refer to as “stable” chromatin sites. Similar to the dynamic sites, the stable regions are associated with gene regulation during NSPC activation, as genes exclusively harboring constitutively accessible chromatin or also associated with dynamic sites both exhibit greater differential expression between qNSCs and aNSCs compared to genes with closed chromatin. This finding shows that stem cells employ at least two distinct chromatin-level mechanisms to support transcriptional programs that regulate early neurogenesis. In the case of stable chromatin, these sites were located at promoters and enriched with the histone mark H3K4me3 that did not change between quiescent and activated NSPCs. Moreover, stable sites were enriched for a distinct set of binding motifs from the dynamic sites. For example, the top candidate transcription factors at stable sites were CTCF and Sp1/5 (zinc finger family transcription factors).

Enrichment of CTCF binding in stable chromatin from activated NSPCs is consistent with our understanding of chromatin domain insulation, and suggests that stable chromatin plays a role in establishing boundaries for gene regulation. Sp1 is a well-known transcriptional activator that binds the proximal promoter. In addition to its ubiquitous expression, Sp1 has also been shown to respond to cellular signals to promote cell proliferation and cell-type specific gene regulation^74, 75^. It will be interesting to investigate the extent to which these factors are already bound at readily accessible sites in quiescent NSCs or are recruited to open chromatin regions during activation to more precisely define the mechanism underlying NSC activation. Nevertheless, our results suggest that in stable chromatin, regulation of gene expression during NSC activation is likely modulated by mechanisms other than “switching on/off” through pioneer factor binding and subsequent alterations in chromatin accessibility, as seen in distal enhancers. Instead, stable sites at active promoters may rely on recruitment of cofactors or a change in the composition of the basal transcriptional machinery to alter gene expression levels^76^. Furthermore, enrichment of the TATA-box and TCT motifs in promoters with stable chromatin identifies a subset of ribosomal genes which are among the most significantly upregulated in NSC activation. To date, it is established that the core promoter is a major contributor to transcriptional regulation^77–79^. One such example is the TCT element that was found to be enriched among ribosomal protein genes, and to regulate their transcriptional output^80^. Recent evidence supports the existence of specialized transcriptional programs through specific transcription factor-core element interactions, in which the distinct core promoter elements serve as docking sites for various basal transcription machineries^77, 81–83^. Our findings suggest that constitutive accessibility of promoters not only allows maintenance of stem cell homeostasis during the transition to activation, but also for regulating particular cellular functions through specific core promoter elements. Taken together, these findings further support the existence of specialized transcription systems that regulate specific biological processes as a function of their core promoter composition, thus providing another dimension to transcriptional regulation. As our understanding of core promoter biology expands, it will be important to probe how each element may function to precisely coordinate specific gene regulation at gene promoters.

NSC quiescence and activation are marked by distinct transcriptional states, giving us a glimpse of the molecular processes that maintain neurogenic potential in the adult brain. Here, for the first time, we identify the chromatin-level changes that accompany the transcriptional overhaul between quiescent and activated NSCs, providing an in-depth understanding of the mechanisms driving NSC activation. This study also raises important questions for future work such as how small and large scale chromatin changes are driven by a combination of pioneer transcription factor activity and chromatin remodeling enzymes to structurally alter chromatin states that fully support activation and functional maturation of NSCs during development and in the adult.

## Methods

### Mouse NSPC cultures

Postnatal (day 5) mouse NSPCs were isolated as previously described^69^. Briefly, whole-mouse forebrains were homogenized and incubated for 30 minutes in HBSS (Invitrogen) with 1 U/ml Dispase II (ThermoFisher), 250 U/ml DNaseI (Qiagen), and 2.5 U/ml Papain (Worthington) at 37°C. After mechanical dissociation, cells were purified by sequential 25% and 65% Percoll (Fisher Scientific) gradients. Cells were cultured in high growth factor signaling conditions: Neurobasal A (ThermoFisher) medium supplemented with penicillin/streptomycin/glutamine (ThermoFisher), 2% B27 (ThermoFisher), and 20 ng/ml each of FGF2 (PeproTech) and EGF (PeproTech). To induce quiescence, 50,000 cells were first seeded in poly-D-lysine coated (Sigma) plates with high growth factor signaling medium, and after 24 hours, fresh Neurobasal A medium with penicillin/streptomycin/glutamine, 2% B27, 20 ng/ml of FGF2, and 25ng/ml of recombinant mouse BMP4 (R&D Systems) was added (without EGF). For reactivated NSPCs, cells were kept in BMP4-containing Neurobasal A medium for 4 days, then switched to high growth factor signaling conditions for 9-12 days. Proliferation of the reactivated NSPCs was confirmed by 10uM EdU (Sigma) incorporation for 2 hours and detection (Click-iT EdU Alexa Fluor 488 Imaging Kit, Invitrogen).

### Immunocytochemistry

NSPCs were plated at 5 × 10^4^ cells/ml on poly-D-lysine (Sigma) treated coverslips. Cells were fixed with 4% paraformaldehyde for 10-15 minutes, blocked for 1 hour with 5% goat serum/0.1% BSA, followed by incubation for 2 hours at room temperature with primary antibody (SOX2 [1:200, EMD Millipore AB5603], NESTIN [1:200, BD Pharmingen #556309]). After washing five times with PBS/0.05% Tween-20, coverslips were incubated with the appropriate secondary antibody for 1 hour at room temperature (Molecular Probes Alexa Fluor, goat anti-rabbit 488, goat anti-mouse 546). Cells were imaged with a Zeiss Axiovert 200M Fluorescence microscope.

### ATAC-seq

ATAC-seq libraries were generated from early passage (passage 2-4) NSPCs cultured in high growth factor signaling conditions (activated), quiescent conditions (quiescent), or high growth factor signaling conditions after quiescence induction (reactivated), with two biologically independent replicates per condition. Library preparations and quality analyses were performed as described^38^. Briefly, for activated NSPCs, 50,000 cells were seeded in Poly-D-Lysine coated plates in high growth factor signaling conditions as described above, and collected with Trypsin-EDTA (ThermoFisher) after 24 hours. For quiescent NSPCs, 50,000 cells were seeded in Poly-D-Lysine coated plates in high growth factor signaling conditions, and after 24 hours, switched to quiescence medium containing FGF2 and BMP4 as described above (no EGF). After 72 hours in quiescence medium, cells were collected with Trypsin-EDTA. For reactivated NSPCs, 50,000 cells were seeded in Poly-D-Lysine coated plates in high growth factor signaling conditions, switched to quiescence medium after 24 hours, and finally switched back to high growth factor signaling conditions after 72 hours for 9 to 12 days. Activated, quiescent, and reactivated NSPCs were subjected to tagmentation reactions with 2.5ul Tn5 Transposase (Illumina), purified with Qiagen MinElute PCR purification kit (Qiagen) and PCR-amplified with 8-9 cycles. Quality of ATAC-seq libraries was confirmed with Bioanalyzer (Agilent) prior to sequencing.

### Processing of ATAC-seq data

Activated and quiescent NSPC ATAC-seq libraries were sequenced to a depth of approximately 40 million unique, high quality mapped reads per sample. 2 × 100 base pair paired-end reads were trimmed with TrimGalore! (Version 0.4.0, Babraham Bioinformatics), and aligned to the most recent genomic builder for *Mus musculus* (Version mm10 from Genome Reference Consortium GRCm38) using Bowtie2 (Version 2.2.5, https://bowtie-bio.sourceforge.net/bowtie2/index.shtml)^84^. Duplicate reads were marked with Picard (Version 1.88, https://broadinstitute.github.io/picard) and removed with SAMtools (Version 1.3.1, https://samtools.sourceforge.net). Reactivated NSPC ATAC-seq libraries were sequenced to a depth of approximately 65 million unique, high quality mapped reads per sample. After processing the reads as described above, reactivated NSPC libraries were downsampled to 40 million reads. Peak calling for all libraries was performed after ATAC-seq specific quality control steps^85^ using MACS (Version 2.1.1)^86^ with the FDR threshold 0.05. Peaks were assigned to genes using GREAT (Version 3.0.0, https://great.stanford.edu/public/html)^87^, limiting the peak-calling window to −1 kb/+1 kb around transcription start sites (TSS) for proximal gene assignments, and −25 kb/+10 kb around TSS for distal gene assignments. For ATAC-seq datasets from postnatal mouse liver, lung, stomach, kidney, and intestine, we downloaded the fastq files from the ENCODE portal^42^ (https://www.encodeproject.org/) with the following identifiers: ENCSR609OHJ, ENCSR102NGD, ENCSR597BGP, ENCSR389CLN, ENCSR079GOY.

### Differential accessibility analysis

The DiffBind package in R was used (Version 3.3.1, https://bioconductor.org/packages/release/bioc/html/DifBind.html)^41^ to identify overlapping and unique ATAC-seq peaks between and across quiescent and activated replicates. DiffBind was used to obtain Pearson’s R correlation values between biological replicates in activated, quiescent, and reactivated conditions, the corresponding heatmap, as well as differential accessibility analysis. Briefly, four peaksets were used as input from which DiffBind derived a “consensus peakset” by merging all overlapping peaks and producing a single set of unique peaks. Next, DiffBind used the sequence reads (BAM) files to count the number of reads that overlap within each peak in the consensus peakset and generate normalized numbers of reads for each sample at every potential site with accessible chromatin. Finally, “dba.report” function was used to generate a list of genomic intervals that were determined to be differentially accessible between quiescent and activated conditions. To identify the peaks in reactivated NSPCs that overlap with dynamic and stable chromatin regions identified in quiescent and activated NSPCs, two peaksets from reactivated NSPC ATAC-seq were used as input from which DiffBind derived a “consensus peakset.” Bedtools (Version 2.26.0) was used to identify the overlaps.

### Genomic distribution of ATAC-seq signals

To determine the distribution of open chromatin regions from ATAC-seq, the ChIPseeker package (Version 3.7, https://bioconductor.org/packages/release/bioc/html/ChIPseeker.html) in R was used^88^. BED files for quiescent and activated ATAC-seq replicates were merged, and promoter regions were defined as −1kb/+1kb around TSSs. Peaks were annotated using “annotatePeak()” command, and pie charts were generated using “plotAnnoPie()”.

### Chromatin Immunoprecipitation

H3K4me3 ChIP-seq libraries were generated from 10 × 10^6^ NSPCs cultured in high growth factor signaling conditions of quiescent conditions. Chromatin was crosslinked with 1% formaldehyde for 10 minutes, followed by quenching with 0.125 M glycine for 5 minutes. Cells were washed with PBS pH 7.4, and incubated in SDS lysis buffer (50 mM Tris-HCl pH 7.5, 10 uM EDTA, 1% SDS) for 15 minutes on ice, and harvested by scraping. Nuclei were pelleted and resuspended in RIPA buffer (1% IGEPAL CA-630, 0.5% sodium deoxycholate, 1% SDS in PBS pH 7.4), and chromatin was sheared with a Covaris S220 Focused-ultrasonicator, at peak power 140 W, duty factor 10% and 200 Cycles/burst at 4°C. Five μg of H3K4me3 antibody (ab8580) was used per ChIP.

### Processing of ChIP-Seq datasets

Raw sequencing reads (FASTQ files) were processed as previously described^89^ with minor modifications for alignment (reference genome mm10 was used instead of mm9). For publicly available datasets, FASTQ files were downloaded from ArrayExpress and Gene Expression Omnibus repository. Briefly, after mapping reads with Bowtie2, duplicate reads were marked with Picard and removed with SAMtools. MACS was used to call peaks using the following commands: “macs2 -t ChIP.bam -c Input.bam -f BAM -g mm -B -q 0.001 --nomodel --extsize 150 --keepdup all”. For the CTCF ChIP-seq dataset, q-value of 1E-8 was used as the cutoff based on the published methods. For the H3K4me3 and H3K27ac ChIP-Seq datasets, an additional parameter of “--broad” was used with “-q 0.05 --broad-cutoff 0.01”. Correlation plots for H3K4me3 ChIP-seq signals between quiescent and activated NSPCs was generated by deepTools (Version 3.1.2, https://deeptools.readthedocs.io/en/develop/)^90^. Peaks were assigned to genes using GREAT, limiting the peak-calling window to −2 kb/+2 kb around transcription start sites (TSS). H3K27ac and p300 binding enrichment around dynamic chromatin regions was performed using NGS plot (https://github.com/shenlab-sinai/ngsplot)^91^.

ChIP-seq datasets downloaded: E-MTAB-1423 (H3K27ac, p300 quiescent and activated NS5 cells, NFI quiescent NS5 cells), GSE48336 (ASCL1), GSE29184 (H3K4me3 in adult and embryonic mouse tissues), GSE85185 (CTCF from postnatal day 1 NSPCs).

### RNA-seq datasets

Generation and analysis of RNA-seq datasets from quiescent and activated in vivo NSCs (freshly isolated from 3-4 month old mouse SVZ) and BMP4-treated and proliferating embryonic stem cell-derived NS5 cells were described previously (Sequence Read Archive accession number SRP075993 for in vivo dataset)^19, 24^. Briefly, for in vivo NSCs, transcript quantification was done using Kallisto^92^. Differential expression of genes was determined by DEseq2^93^, normalizing the read counts using the variance stabilizing transformation (VST). For NS5 cells, transcripts were processed with TopHat and the Cufflinks package^94^. For this study, published normalized expression and fold change values were used.

### Violin plots and statistical analysis of differential expression

Violin plots were created in R using ‘ggplot2’, with the ‘geom_violin,’ and geom_boxplot’ functions (http://ggplot2.org/). Statistical analysis of the gene expression differences between quiescent and activated NSPCs, with and without dynamic chromatin, was performed using Wilcoxon signed rank sum test with continuity correction.

### Venn diagrams and statistical analysis of overlaps

Venn diagrams were created in R using the ‘venneuler’ package (https://www.rforge.net/venneuler). Statistical analysis was performed using Fisher’s exact test. When calculating significance, we included genes without changes in expression between quiescent and activated conditions in addition to the differentially expressed genes^24^ for the ‘background dataset.’

### Lentiviral constructs and infection

gRNAs targeting a putative enhancer region upstream of *Aqp4* (chr18:15421139-15421834) were generated using GT-Scan^95^ and selected based on minimal number of off-targets. gRNAs were cloned into the lentiviral gRNA expression plasmid, pLV-U6-gRNA-UbC-DsRed-P2A-Bsr (Addgene plasmid #83919), by Gibson assembly as previously described^50, 96^. See Supplementary Table 7 for oligonucleotide sequences. pLV-dCas9-KRAB-PGK-HygR (Addgene plasmid #83890) was described previously^50^. Lentiviruses were produced in HEK293T cells using polyethylenimine (PEI) (Polysciences)^97^ transfection including the packing plasmids PMDL, VSV-G, RSV. After 24 hours, we replaced the transfection media with fresh Neurobasal A medium with penicillin/streptomycin/glutamine and 2% B27. Lentiviral media was collected 24 hours later. To generate dCas9-KRAB-expressing cells, NSPCs were plated on poly-D-lysine at 50,000 cells/cm^2^ and infected with 1:1 ratio of growth media to viral supernatant. Virus was removed after 24 hours and replaced with growth media. 3 days after infection, cells were selected with 700 ug/ml Hygromycin (Fisher Scientific) for 6 days. dCas9-KRAB-expressing NSPCs were induced into quiescence with BMP4 as described above, and simultaneously infected with 1:1 ratio of growth media to gRNA lentiviral supernatant. Virus was removed after 24 hours, and infection was confirmed with DsRed fluorescence. RNA was collected 7 days after infection.

### RT-qPCR

Total RNA was isolated using the QIAGEN RNeasy kit with on column DNase digestion (QIAGEN). 250 – 500 ng RNA was used for reverse transcription, which was performed using the High Capacity cDNA Reverse Transcription Kit (Applied Biosystems). qPCRs were performed using Powerup SybrGreen Master Mix (Invitrogen) and run on an Applied Biosystems ViiA 7 Real-Time PCR System. See Supplementary Table 7 for primer sequences used.

### Motif analysis

Motif analysis was performed using the Homer findGenomeMotifs.pl tool (http://homer.salk.edu/homer/index.html)^98^ on dynamic and stable chromatin regions between activated and quiescent NSPCs. Default settings (200 base pair window) were used in all analyses, and *P-*values from the Homer Known Motif Enrichment Results are reported. In cases where a motif appeared multiple times, only the top (smallest) *P*-value is reported.

### ElemeNT analysis

Core promoter analysis was performed using ElemeNT (Elements Navigation Tool)^48^ command line version (version 13). Constitutively accessible promoters of upregulated genes in NSC activation in vivo were divided into quartiles by rank (FDR for differential expression). To detect element motifs, −50/+50 bp window around TSS that is stably open was analyzed. For each element, the positions from the input start position and corresponding PWM scores were reported. For enrichment analysis, −50/+50 bp window around TSS for all constitutively accessible gene promoters was used as background.

Enrichment significance was calculated using the hypergeometric distribution and adjusted by Bonferroni correction for multiple testing. Only genes with element motifs in biologically relevant positions (-30/-31 to −23/-24 from TSS for the TATA box element, −1 to +6 from the TSS for the TCT motif, with −10/+10 bp allowance around expected positions) were highlighted in Fig. 4e.

### Functional annotation

Functional annotation of genes that are associated with dynamic chromatin regions and/or differential expression with NSC activation was performed using Ingenuity Pathway Analysis (Qiagen Inc., https://www.qiagenbioinformatics.com/products/ingenuity-pathway-analysis)^46^. For IPA core analysis, tables of genes and their differential expression in vivo were uploaded. Enriched canonical pathways were plotted in R using ‘ggplot2’.

## Code availability

The code will be released at: https://www.thewebblab.com.

## Data availability

The ATAC-seq and ChIP-seq data from this study will be deposited in NCBI’s Gene Expression Omnibus (GEO).

## Author contributions

S.M-L. and A.E.W. conceived and designed the study. S.M-L. and S.D. harvested primary mouse NSPCs for experiments. S.M-L. performed ATAC-seq and ChIP-seq with the supervision of A.E.W. S.M-L. performed processing and analysis of ATAC-seq and ChIP-seq data to investigate epigenomic changes between quiescent and activated NSPCs. A.K.B. performed immunocytochemistry experiments to validate the neural stem cell identity in NSPCs. M.Y. set up the dCas9-KRAB system. B.M-S. performed RT-qPCR experiments on candidate transcriptional regulators from Homer binding motif analysis. All remaining experiments were done by S.M-L. A.S. and T.J-G provided guidance on core promoter analysis, as well as helpful comments on the manuscript. S.M-L and A.E.W. prepared the manuscript.

## Acknowledgements

We thank members of the Webb laboratory for critical reading of the manuscript. This work was supported by NIH training grant 5T32AG041688, Glenn/AFAR Scholarship for Research on the Biology of Aging, and Brown University Carney Brain Institute for Brain Science Robin Chemers Neustein Graduate Research Award to S.M-L., as well as NIH grant R01 AG053288 to A.E.W.

## Supporting Information

### Supplementary tables

**Supplementary Table 1.** Read numbers for ATAC-seq libraries and peak calling results.

**Supplementary Table 2.** Summary of DiffBind differential accessibility analysis.

**Supplementary Table 3.** Summary of GREAT gene assignments to dynamic and stable chromatin sites.

**Supplementary Table 4.** Read numbers for H3K4me3 ChIP-seq libraries and peak calling results.

**Supplementary Table 5.** Complete tables of Ingenuity Pathway Analysis (IPA) results.

**Supplementary Table 6.** Complete tables of core promoter element analysis (ElemeNT) results.

**Supplementary Table 7.** Oligonucleotide sequences for lentivirus and RT-qPCR experiments.

### Supplementary figures

**Supplementary Fig. 1.**
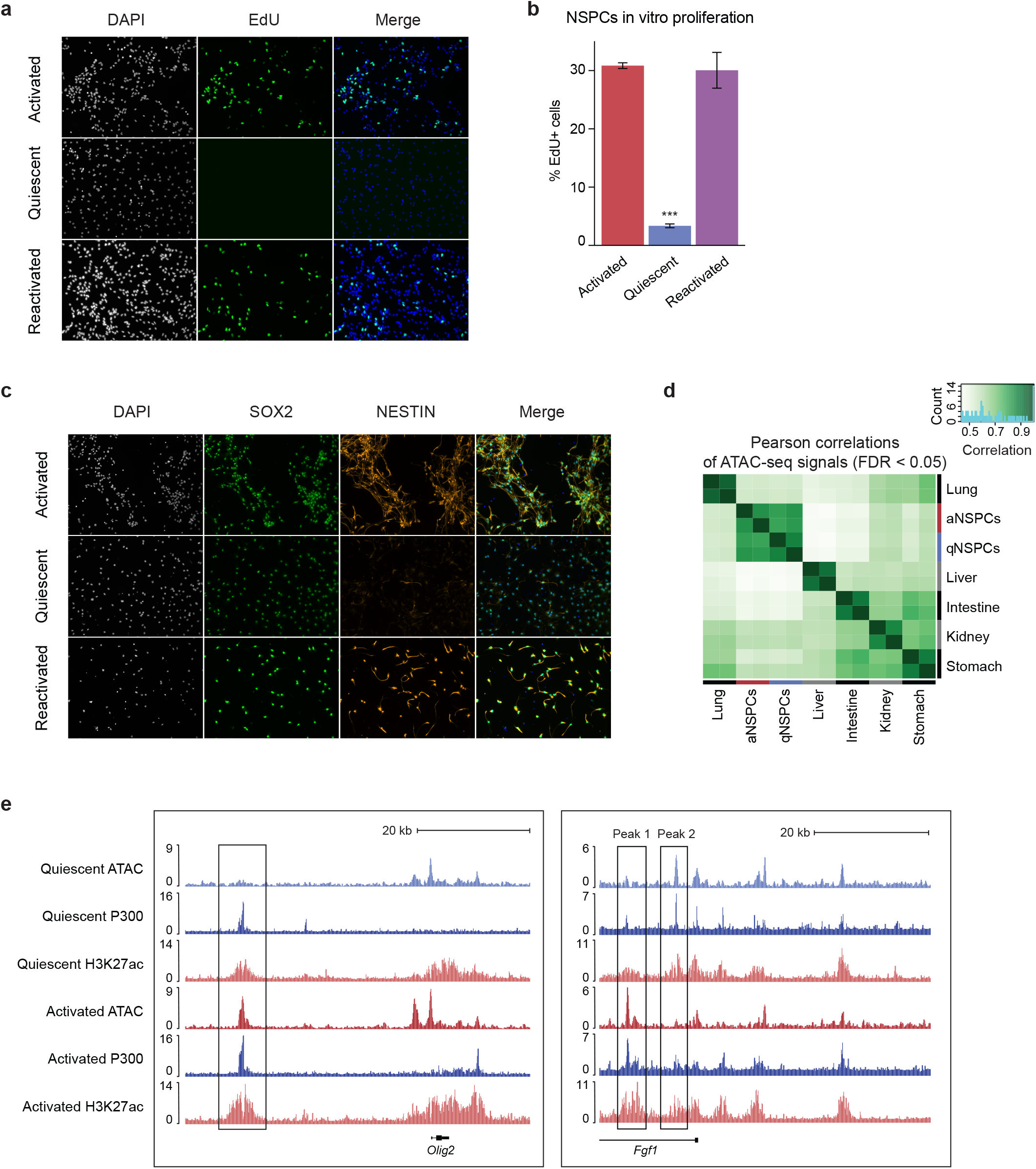
Activated and BMP4-induced quiescent NSPCs express NSC markers and are distinct from other cell types in their open chromatin profiles. (a) Representative images of EdU-positive NSPCs in activated, quiescent, and reactivated states. (b) Percentage of EdU-positive cells in activated, quiescent, and reactivated states. (n = 3, Student’s t-test, \*\*\**P* < 0.001). (c) SOX2 and NESTIN expression in activated, quiescent, and reactivated NSPCs. Green, SOX2; orange, NESTIN. (d) Heatmap depicting Pearson correlation of ATAC-seq signals among quiescent and activated NSPCs and other mouse tissues (lung, liver, intestine, kidney, stomach). (e) UCSC genome browser snapshots of *Olig2* (left) and *Fgf1* (right) locus with quiescent and activated ATAC-seq, H3K27ac and p300 ChIP-seq tracks marking active enhancers.

**Supplementary Fig. 2.**
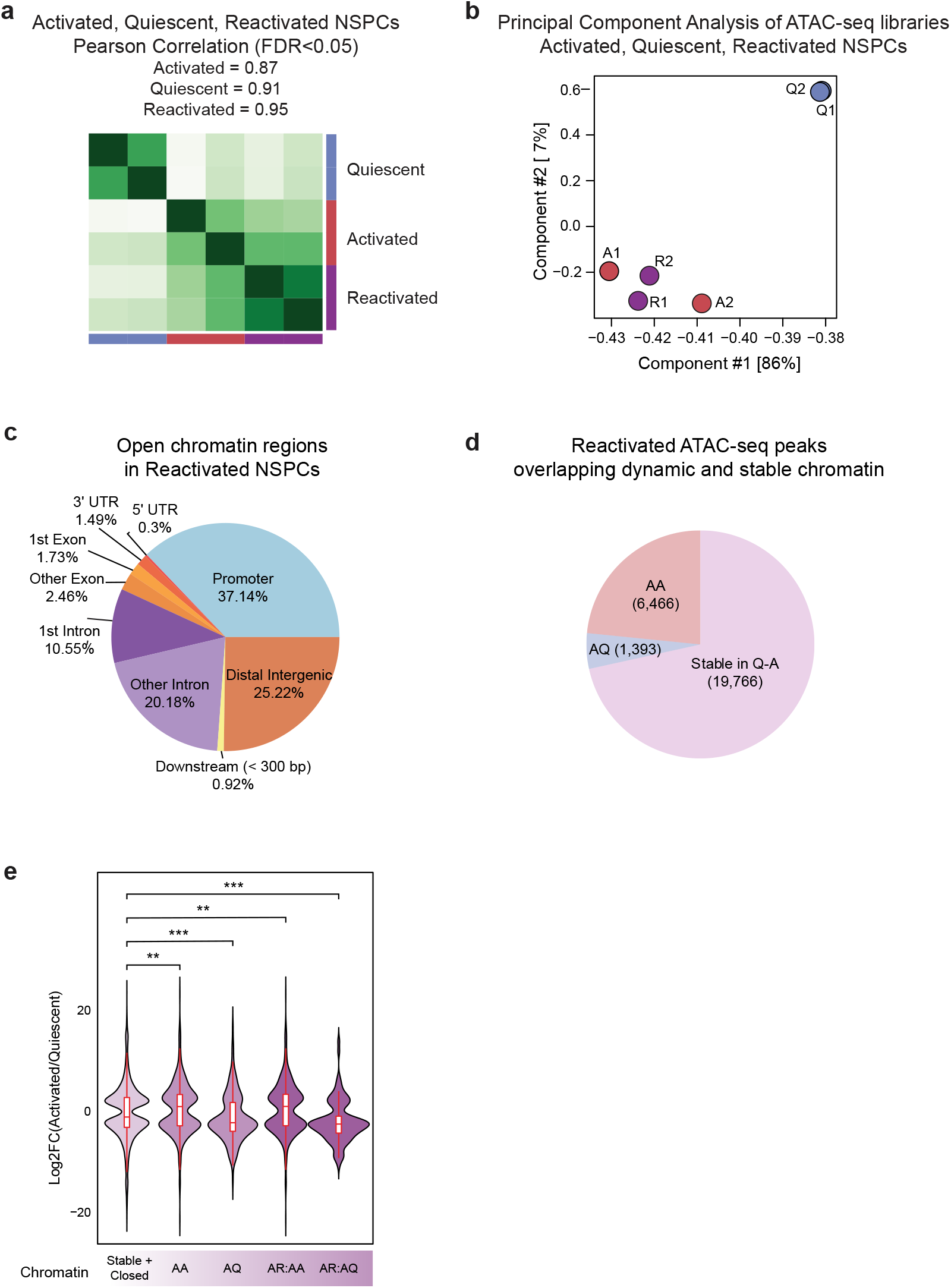
The chromatin accessibility profile of reactivated NSPCs resembles that of activated NSPCs. (a) Heatmap depicting Pearson correlation coefficient r (0.87 in activated, 0.91 in quiescent, 0.95 in reactivated) in ATAC-seq signals between biological replicates in activated, quiescent, and reactivated conditions. (b) Principal component analysis of biological replicates in activated, quiescent, and reactivated ATAC-seq. (c) Genomic distribution of all open chromatin regions in reactivated NSPCs. (d) Pie chart depicting the percentages of overlaps among open chromatin regions in reactivated NSPCs and stable or dynamic (AA or AQ) chromatin regions in quiescent and activated NSPCs. (e) Comparison of differential expression in vivo (fold change) between genes with chromatin that does not change, AA chromatin, AQ chromatin, AA chromatin accessible in reactivated NSPCs (AR:AA), and AQ chromatin accessible in reactivated NSPCs (AR:AQ) (Wilcoxon signed rank test, \*\*\**P* < 0.001, \*\**P* < 0.01).

**Supplementary Fig. 3.**
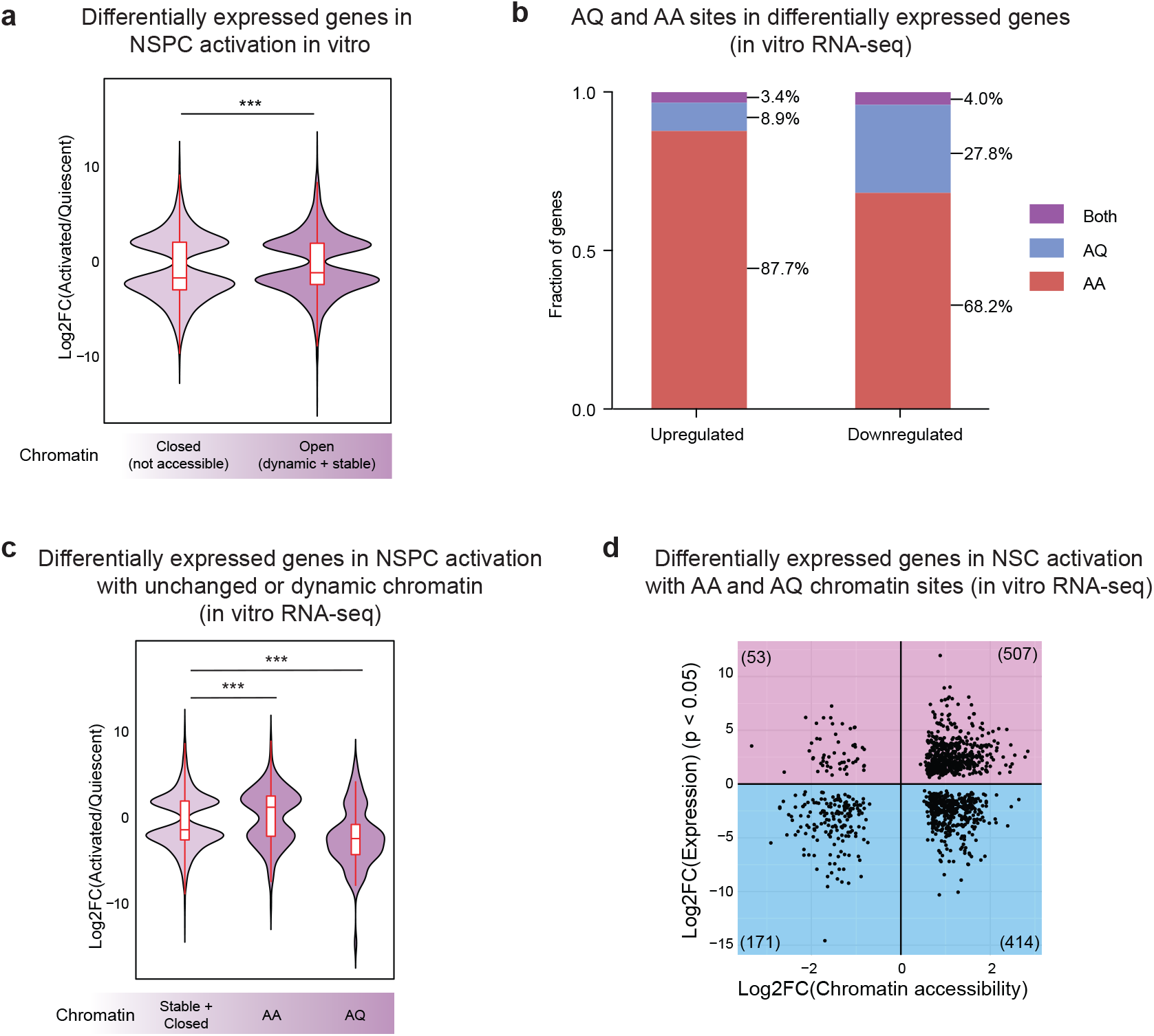
Dynamic chromatin regions are associated with differential expression of genes between quiescent and activated states in vitro. (a) Comparison of differential expression (fold change) between genes with closed or open (stable and dynamic) chromatin. RNA-seq data from cultured NS5 cells show that genes with open chromatin are more differentially expressed in vitro compared to genes with closed chromatin (Wilcoxon signed rank test, *\*\*\*P* < 0.001). (b) Bar graph depicting percentages of differentially expressed genes in vitro that contain dynamic (AQ or AA) chromatin sites, or both. (c) Comparison of differential expression of genes with chromatin showing no change (constitutively accessible and closed chromatin) and AA and AQ dynamic chromatin (Wilcoxon signed rank test, \*\*\**P* < 0.001). (d) Scatterplot depicting fold change in chromatin accessibility versus fold change in associated gene expression in vitro (FDR < 0.05). Each dot represents a dynamic site that is associated with a differentially expressed gene, and the total number of sites in each quadrant is shown in parentheses. Downregulated genes are enriched in AQ sites (Fisher’s exact test, *P* = 3.2 × 10^−12^).

**Supplementary Fig. 4.**
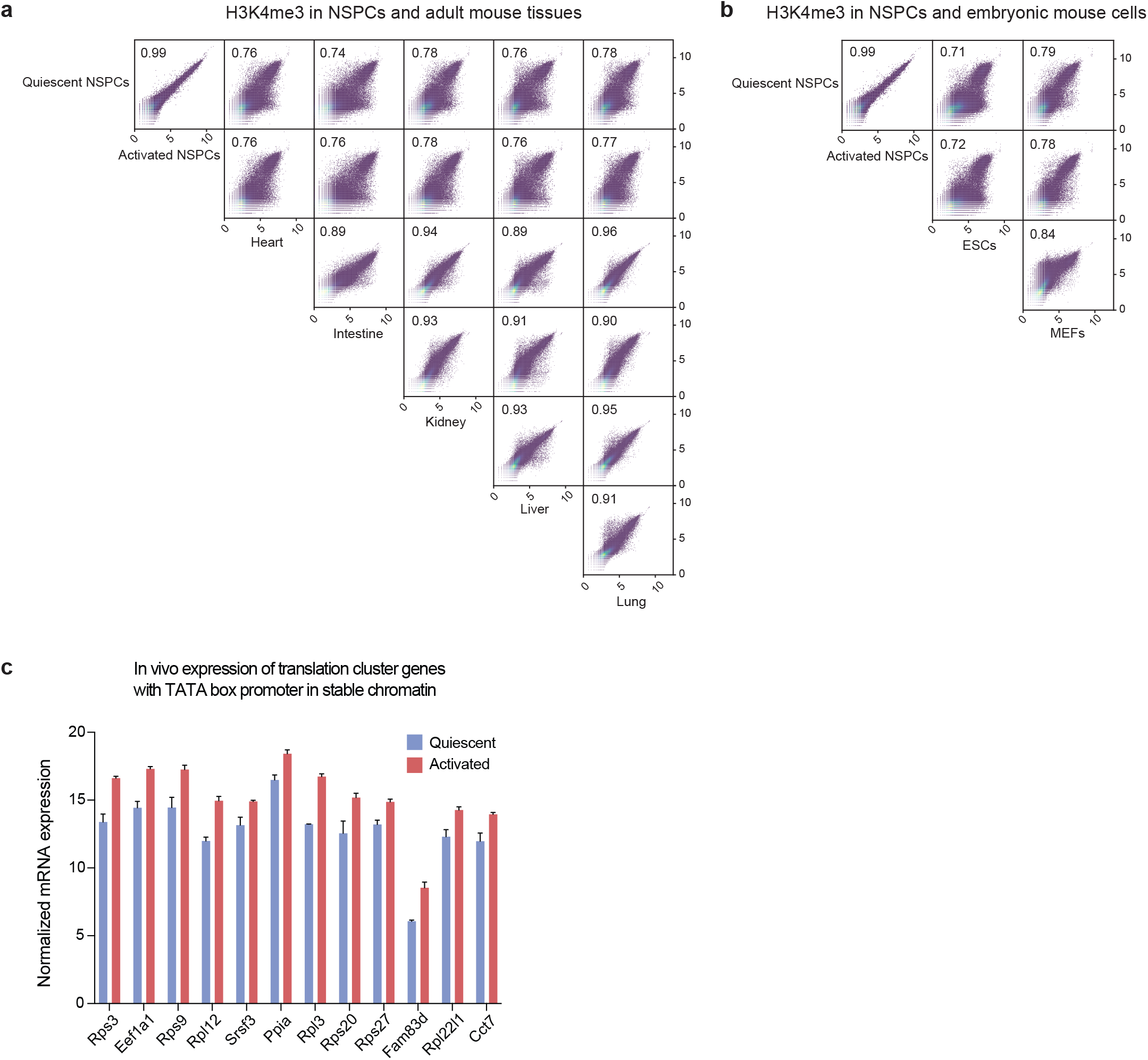
Constitutively accessible chromatin regions in NSPCs are enriched with cell type-specific active promoter marks and specific core promoter elements. (a) Heatmap depicting Pearson correlation coefficient r among ATAC-seq signals from quiescent and activated NSPCs and published 8-week-old adult mouse tissues^47^. Correlation coefficients are shown in upper left corner of each plot. (b) Heatmap depicting Pearson correlation coefficients among ATAC-seq signals from quiescent and activated NSPCs and embryonic mouse cells (ESCs and MEFs). (c) Normalized mRNA expression values from quiescent and activated in vivo NSCs. A cluster of ribosomal and translation-associated genes are consistently upregulated in NSC activation, and harbor the TATA box motif in their core promoters (-40/-41 to −13/-14 bp from TSS).

**Supplementary Fig. 5.**
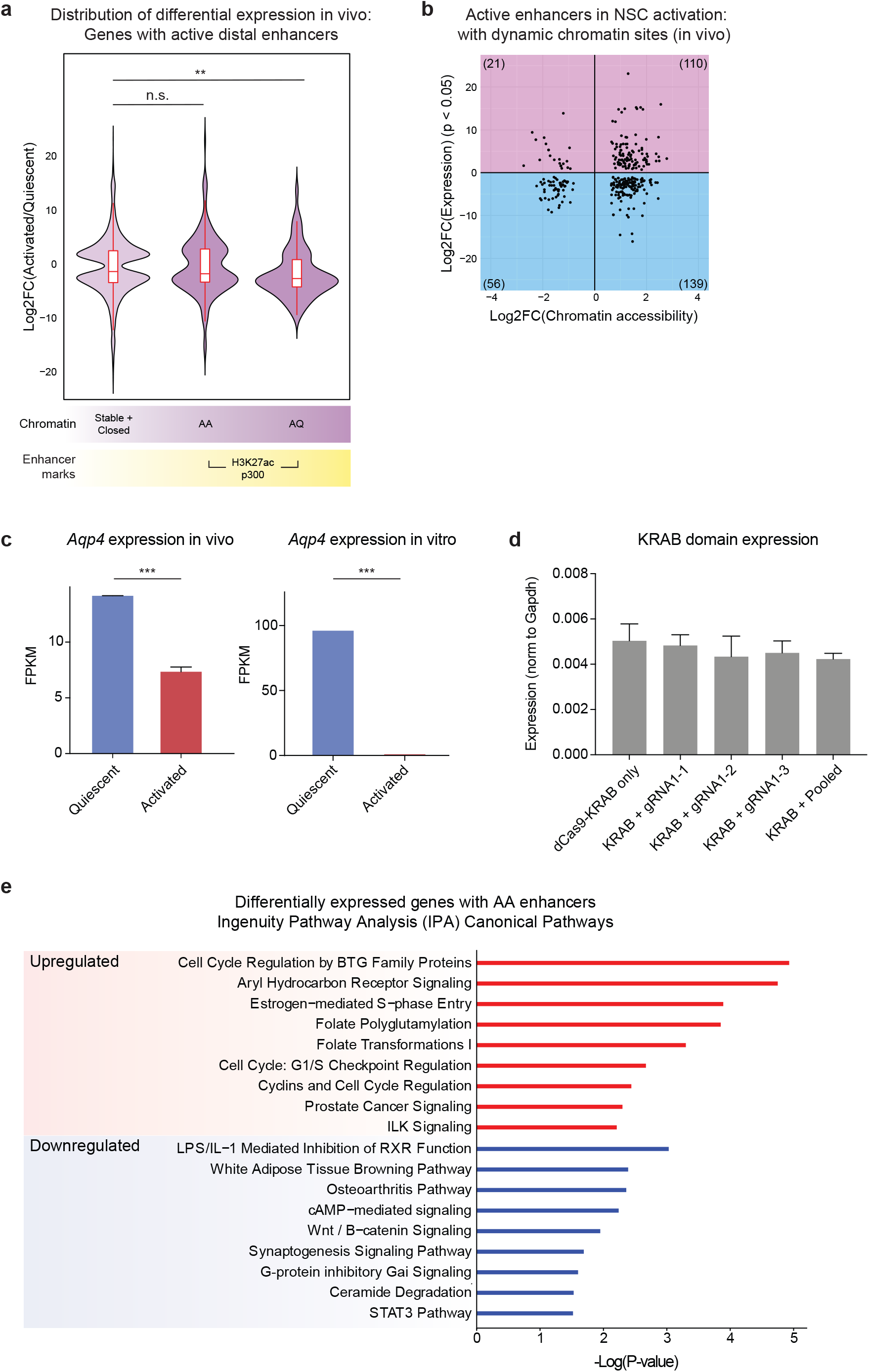
Active enhancers in dynamic chromatin and their association to differential gene expression. (a) Comparison of differential expression in vivo between genes with constitutively accessible or closed chromatin and genes with active enhancers in dynamic chromatin (Wilcoxon signed rank test, \*\**P* < 0.01). (b) Scatterplot showing fold change in chromatin accessibility versus fold change in associated gene expression (FDR < 0.05). Each dot represents a dynamic chromatin site with active enhancer marks, and is associated with a differentially expressed gene. Total number of sites in each quadrant is shown in parentheses. (c) Differential expression of *Aqp4* in quiescent and activated NSCs in vivo (left) and cultured NS5 cells (right). *Aqp4* is significantly upregulated in quiescent cells (FDR-corrected \*\*\**P* < 0.001). (d) RT-qPCR analysis of the *KRAB* repressor domain in NSPCs infected with lentiviruses carrying dCas9-KRAB and gRNAs (n = 3 experiments). (e) Ingenuity Pathway Analysis (IPA) Canonical Pathways of genes differentially expressed in vivo and associated with AA enhancers. Top 9 pathways are shown for genes upregulated and downregulated with NSC activation.

**Supplementary Fig. 6.**
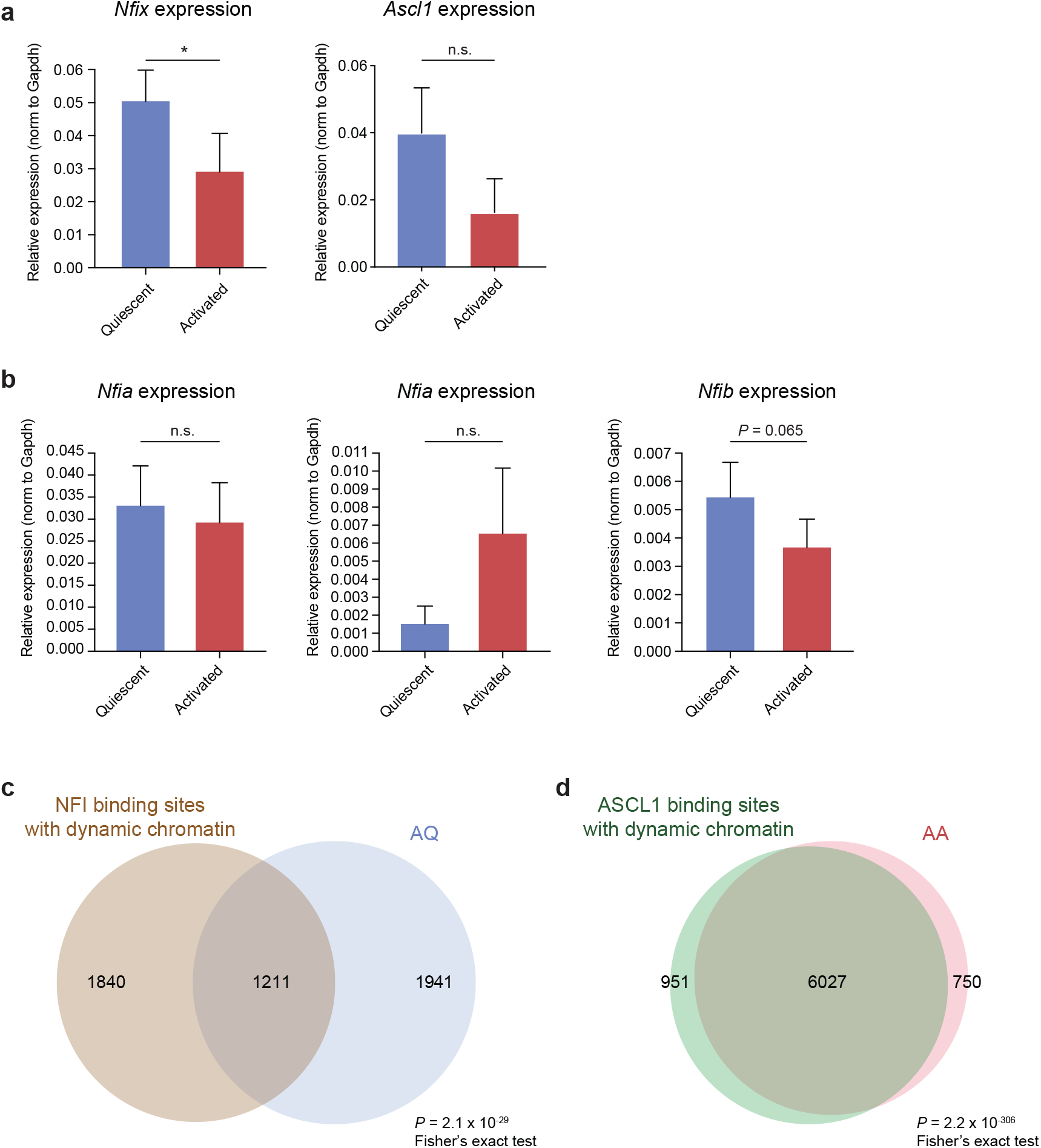
Dynamic accessible chromatin regions are enriched for binding of ASCL1 and NFI factors. (a) RT-qPCR analysis of *Nfix* and *Ascl1* expression in quiescent and activated NSPCs (n = 3, Welch’s t-test, \**P* < 0.05). (b) RT-qPCR analysis of *Nfia*, *Nfib*, and *Nfic* expression in quiescent and activated NSPCs (n = 3, Welch’s t-test). (c) Overlap of NFI binding with AQ chromatin sites (Fisher’s exact test, *P* = 2.1 × 10^−29^). (d) Overlap of ASCL1 binding with AA chromatin sites (Fisher’s exact test, *P* = 2.2 × 10^−306^).

